# Rebound Relays and Inhibitory Vetoes Stabilize Sparse Sequential Activity in HVC

**DOI:** 10.64898/2026.03.07.710036

**Authors:** Zeina Bou Diab, Arij Daou

## Abstract

Brains build behavior by chaining actions and perceptions into precisely timed sequences, an ability central to speech, skilled movement, and memory, yet the circuit logic that propagates sequences remains unclear. Songbird HVC captures this problem: premotor HVC_RA_ neurons burst once per motif, while basal ganglia-projecting HVC_X_ neurons burst 2-4 times across the same motif. We developed a biophysically grounded HVC network as linked microcircuits encoding sub-syllabic segments. The model highlights inhibition not merely as suppressive but actively structuring sequence propagation and fidelity. Tonic inhibitory epochs prime HVC_X_ by deinactivating T-type Ca^2+^ channels and recruiting *I*_*h*_, so that release elicits precisely timed rebound bursts that recruit the next HVC_RA_ ensemble. A complementary phasic inhibitory veto suppresses off-time activation, preventing pathological restarts while preserving HVC_RA_ single-burst sparseness. More broadly, inhibitory timing can serve as the brain’s internal “clocked handshake,” converting suppression into forward drive to advance sequences while enforcing error-corrected precision.

## Introduction

The ability to learn and generate ordered patterns of activity is a cornerstone of cognition and behavior. Across systems - navigation, decision making, and skilled movement - neural populations represent time not as a static variable but as structured trajectories that enable prediction, planning, and action^1–6^. Yet a mechanistic question remains central: how do circuits propagate precise sequences while preventing spurious reactivation and preserving cell-type specific firing patterns?

Songbird vocal behavior provides a canonical model for addressing this question in a learned sensorimotor sequence. Birds acquire song through iterative practice guided by auditory feedback, and adulthood retains flexibility for ongoing maintenance and modification. This learning is supported by a well-defined circuit architecture comprising a vocal motor pathway and a cortico-basal ganglia loop (Figure 1 in Bou Diab et al^7^), enabling unusually direct links between circuit dynamics, behavioral output, and plasticity. At the core of this system lies HVC, a telencephalic nucleus often considered homologous to mammalian cortical layer III^8,9^. HVC functions as a pattern generator and “conductor” of song, encoding syllable order and fine temporal structure through stereotyped commands that ultimately drive vocal and respiratory musculature^10–15^.

**Figure 1.**
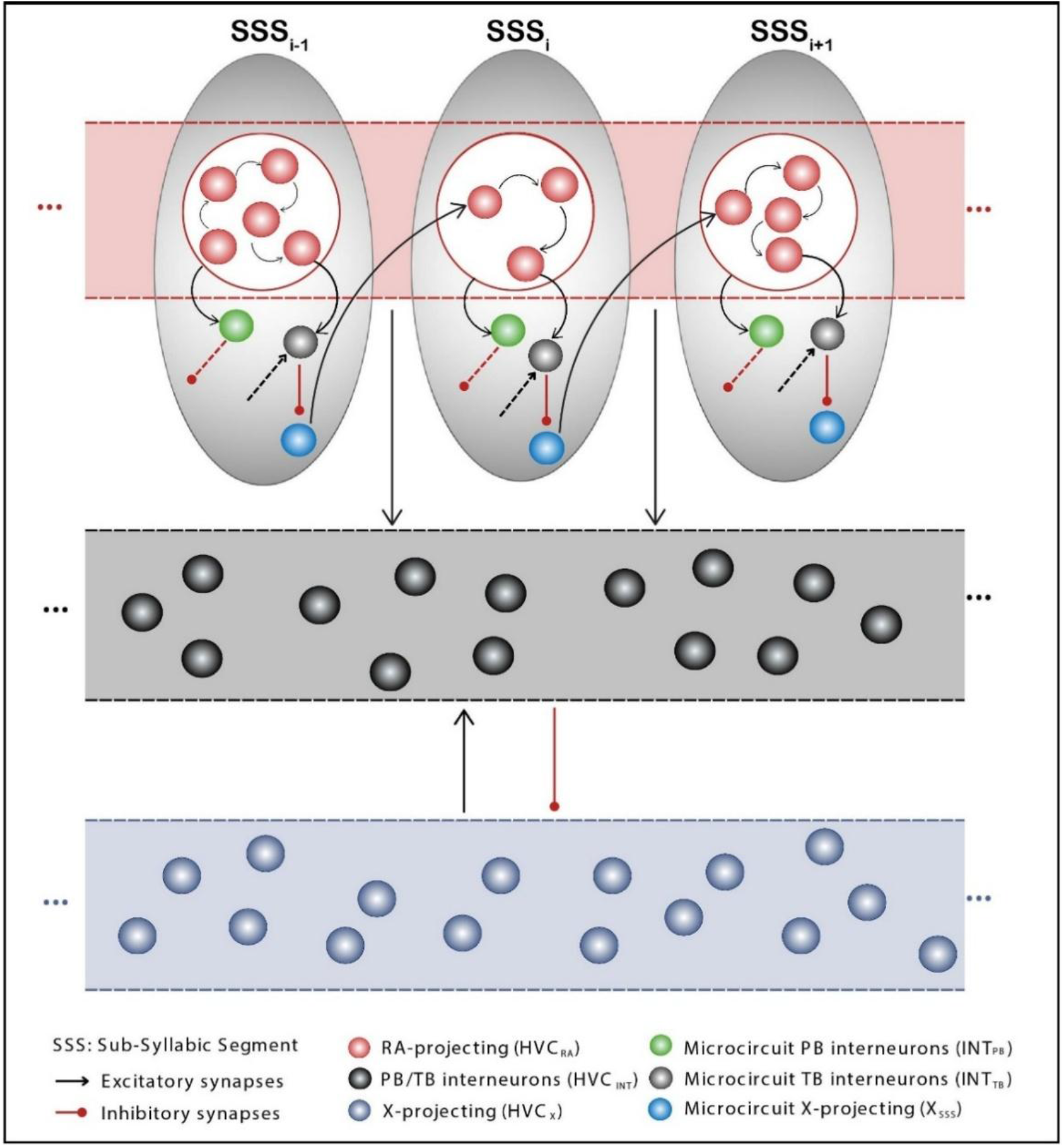
Microcircuit-chain architecture of the HVC network model. HVC is modeled as a chain of repeating microcircuits, each corresponding to one sub-syllabic segment (SSS) of the motif (illustrated for *SSS*_*i*−1_, *SSS*_*i*_, and *SSS*_*i*+1_). Within each microcircuit (gray oval), a small ensemble of RA-projecting neurons (HVC_RA_; red) is connected by directed AMPAergic links (black arrows) and recruits three microcircuit-specific elements: a phasic-bursting interneuron (*INT*_*PB*_; green), a tonic-bursting interneuron (*INT*_*TB*_; gray), and a microcircuit-specific X-projecting neuron (*X*_*SSS*_; dark blue). HVC_RA_ activity drives local inhibition; *INT*_*TB*_ inhibits *X*_*SSS*_, which then provides excitatory drive to the first HVC_RA_ neuron of the next microcircuit, implementing segment-to-segment propagation. Microcircuits are embedded within global pools of non-microcircuit-specific interneurons (PB/TB HVC_INT_; emphasized black zone) and HVC_X_ neurons (emphasized blue zone) that interact sparsely and reciprocally, providing distributed background coupling across segments. Black arrows denote excitatory synapses and red symbols denote inhibitory synapses; ellipses indicate repetition across the motif.

HVC is mechanistically tractable because it comprises a small set of principal cell classes with distinct connectivity and firing regimes. HVC_RA_ neurons provide timing signals to the robust nucleus of the arcopallium (RA) and downstream motor centers^14,16^, while HVC_X_ neurons project to basal ganglia Area X, linking sequence generation to reinforcement-driven learning^17^. Local fast-spiking interneurons (HVC_INT_) densely regulate both projection classes, shaping timing and reliability of activity. *In vivo*, this division is reflected in strikingly different spike patterns: during singing, HVC_RA_ neurons fire a single brief burst (~10 ms) locked to a specific moment in the motif^18,19^, HVC_X_ neurons generate multiple bursts (2-4) that are likewise stereotyped and motor-locked^19,20^, and HVC_INT_ neurons exhibit sustained firing with structured epochs of bursting and suppression^18,19,21–25^. Recent reanalysis further suggests that HVC_X_ bursting may form multiple concurrent motif-spanning sequences (“parallel lines”), pointing to a richer temporal code in HVC_X_ than previously appreciated^26,27^.

Decades of intracellular work constrain the intrinsic and synaptic mechanisms available to produce these patterns. *In vitro* and *in vivo* recordings demonstrate that HVC_RA_, HVC_X_, and HVC_INT_ exhibit distinct electrophysiological signatures shaped by specific complements of ion channels^28–34^. Paired recordings in slices revealed a core microcircuit motif: projection neurons excite interneurons, interneurons inhibit both projection classes, with evidence for monosynaptic excitation from HVC_X_ to HVC_RA_ embedded in a larger landscape of disynaptic inhibition^24,32^. Multiple computational models have been proposed for HVC sequence generation, invoking intrinsic chaining, extrinsic feedback, or both^7,13,23,25,35–42^. Yet most do not simultaneously capture realistic spike morphology with biophysical currents, include all major HVC classes, and reproduce the distinct in vivo firing patterns across populations; even recent biophysically grounded microcircuit models leave open how HVC_X_ and interneurons causally advance sequences and generate motif-spanning HVC_X_ structure^7,27^.

Recent causal and synaptic-mapping work has sharpened key constraints. Trusel et al.^43^ showed that once song is initiated, HVC can sustain progression through the learned motif largely independent of major extrinsic input pathways, and that brief excitation within HVC can trigger “skipping record”-like restarts that rely on local circuitry. Importantly, their connectivity mapping finds HVC_X_ neurons consistently making monosynaptic connections onto HVC_RA_ neurons, but only very sparsely onto other HVC_X_ neurons, supporting a sequence-generating network built around heterotypic excitation between the two projection classes. These results elevate HVC_X_ from a corollary channel for basal ganglia learning to a potential causal relay within the premotor pattern generator itself.

Here we present a biophysically grounded model of HVC that reframes sequence generation around that relay. The model organizes HVC as a chain of linked microcircuits encoding sub-syllabic segments and integrates experimentally constrained synaptic architecture with realistic intrinsic currents. We show how tonic inhibition primes HVC_X_ neurons for post-inhibitory rebound via T-type Ca^2+^ deinactivation and *I*_*h*_ recruitment, so that release generates precisely timed bursts that recruit the next HVC_RA_ ensemble, while a complementary phasic inhibitory veto suppresses off-time reactivation to preserve HVC_RA_ single-burst sparseness. This framework links defined cellular conductances to motif-scale sequence integrity and yields concrete, testable predictions for how manipulating GABAergic tone or HVC_X_ intrinsic currents should reshape propagation, burst multiplicity, and timing.

## Results

Zebra finch song is an exceptionally precise learned sequence: each motif comprises stereotyped syllables produced with millisecond timing across renditions. In HVC, this precision is expressed as distinct population codes; HVC_RA_ neurons emit a single brief burst per motif time point, HVC_X_ neurons generate multiple motif-locked bursts, and local interneurons fire densely with structured epochs of activation and suppression. Dominant synaptic motifs and key intrinsic conductances have been characterized across these classes^24,32,34,44^, providing unusually strong constraints for mechanistic modeling.

Here we build a biophysically grounded HVC network to ask what circuit mechanisms are sufficient to propagate a sparse sequence with HVC_RA_ single-burst precision while preserving multi-burst HVC_X_ dynamics and realistic interneuron activity. We first define a microcircuit-chain architecture (Figure 1), then show how propagation is carried by an HVC_X_-mediated relay that converts inhibition into forward drive, while complementary inhibition suppresses spurious HVC_RA_ reactivation. Finally, we map parameter regimes that stabilize propagation and predict perturbations that fragment the sequence, change HVC_X_ burst multiplicity, or warp motif timing.

### Network Architecture: HVC_X_ to HVC_RA_ driven activity

We modeled HVC as a chain of repeating microcircuits arranged in a functional chain, each representing the minimal unit required to generate activity for a sub-syllabic segment (SSS)^7^. A motif composed of *N* SSSs is implemented as *N* linked microcircuits, each recruiting a dedicated ensemble whose coordinated intrinsic and synaptic dynamics both encode the local segment and advance activity to the next. This framing converts the core problem of song timing into a mechanistic one: identify the minimal, biophysically grounded circuit ingredients that can propagate a sparse sequence while preserving the distinct firing regimes of HVC_RA_, HVC_X_, and interneurons.

#### Structural organization of the network

Figure 1 summarizes the network architecture and population structure. The model contains a total pool of 120 HVC_RA_ neurons (red), 60 HVC_X_ neurons (light/dark blue), and 60 interneurons split into 20 phasic-bursting interneurons (INT_PB_) (green) and 40 tonic-bursting interneurons (INT_TB_) (light/dark black), preserving the reported 2:1:1 proportionality across HVC_RA:_ HVC_INT_: HVC_X_^31,44^. For computational tractability, we instantiated 20 microcircuits (i.e., 20 SSSs); both the number of microcircuits and absolute population sizes are scalable without changing the architectural logic. We use two operational interneuron classes to capture the inhibitory motifs required for realistic output: PB interneurons produce a single, time-locked burst, whereas TB interneurons generate multiple bursts across the motif. As shown below, these modes play complementary roles in propagation and in preventing spurious sequence reactivation.

Each microcircuit (gray oval) contains a **microcircuit-specific** (MS) HVC_RA_ ensemble - a random group of HVC_RA_ neurons drawn from the global pool (ensemble size chosen randomly; see below) - and exactly three MS control neurons: one INT_PB_, one INT_TB_, and one MS HVC_X_ neuron (X_SSS_). “Microcircuit-specific” denotes neurons that belong to one and only one microcircuit and are required for robust sequential propagation. All remaining interneurons and HVC_X_ neurons are non-microcircuit-specific (NMS), and can receive/deliver synapses broadly (subject to pre/post cell-type rules), providing a distributed inhibitory-excitatory background that interacts with microcircuit dynamics (Figure 1).

#### Synaptic connectivity in the network

To translate the microcircuit-chain hypothesis into a mechanistic network, we implemented a connectivity scheme that is minimal, cell-type grounded, and explicitly designed to separate three functions: (i) generate a brief, sparse HVC_RA_ burst packet within each microcircuit, (ii) recruit precisely timed inhibition that shapes burst termination and de-inactivation dynamics, and (iii) transmit activity forward to the next microcircuit through an HVC_X_-mediated relay.

### Feedforward HVC_RA_ chaining within each microcircuit

The 120 HVC_RA_ neurons are partitioned across 20 microcircuits, with each microcircuit recruiting a random HVC_RA_ ensemble of 3-10 neurons. Within microcircuit *i*, these neurons are ordered into a directed AMPAergic chain: 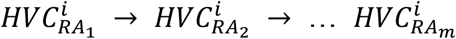 (Figure 1). This implementation enforces a key constraint of the song system: local excitation is temporally ordered, providing a transparent substrate for generating a brief, temporally ordered HVC_RA_ packet.

### Microcircuit-specific inhibition splits into phasic and tonic control

Each microcircuit contains two inhibitory controllers with distinct targets and roles. All HVC_RA_ neurons in microcircuit *i* excite 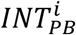 via AMPA synapses, making 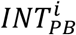 a rapid readout of microcircuit activation positioned to deliver precisely timed inhibition that prevents unwanted reverberation (mechanism dissected below). In contrast, only the terminal HVC_RA_ neuron 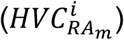 excites 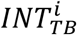, implementing an “end-of-segment” signal that couples TB recruitment to microcircuit completion. In addition, each 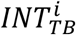 can receive excitation from a random subset of HVC_RA_ and HVC_X_ neurons across the network, and each 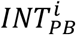 can inhibit selected HVC_RA_ or HVC_X_ (Figure 1).

### HVC_X_ relay converts inhibition into forward drive

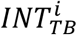 inhibits the MS HVC_X_ neuron 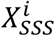 via GABAergic synapses. Critically, 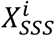 then excites the first HVC_RA_ neuron of the next microcircuit, 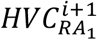, via an AMPA synapse. This single motif - HVC_RA_ chain → TB inhibition → HVC_X_ rebound → next HVC_RA_ chain - is the backbone of propagation in the model, and is consistent with recent evidence for robust HVC_X_ → HVC_RA_ connectivity^43^.

### Distributed NMS coupling

To introduce realistic background interactions without hardwiring additional sequence structure, we included sparse probabilistic coupling through NMS pools. HVC_RA_ neurons provide random AMPA excitation to NMS interneurons (each HVC_RA_ contacting 1-3 randomly chosen NMS interneurons). NMS interneurons inhibit NMS HVC_X_ neurons (1-3 targets per interneuron), and NMS HVC_X_ neurons reciprocally excite NMS interneurons (1-3 targets per HVC_X_). These low-density interactions supply a distributed inhibitory-excitatory backdrop while keeping sequence propagation primarily determined by the microcircuit relay.

### Functional connectivity implements propagation while enforcing cell-type specific firing regimes

Figure 2 summarizes how the model’s two inhibitory modes - phasic and tonic bursting interneurons - coordinate with HVC_RA_ and HVC_X_ neurons to propagate activity across sub-syllabic segments (SSSs) while preserving each class’s *in vivo*-like firing regime. Traces are shown for four example microcircuits (gray boxes): two consecutive segments (microcircuits *i* and *i* + 1, encoding *SSS*_*i*_ and *SSS*_*i*+1_) and two later, nonconsecutive microcircuits (*j* and *k*, encoding *SSS*_*j*_ and *SSS*_*k*_). Each microcircuit recruits a randomly sized HVC_RA_ ensemble (here 5, 3, 6, and 7 neurons, respectively), reflecting stochastic assignment of 3-10 HVC_RA_ neurons per microcircuit.

**Figure 2.**
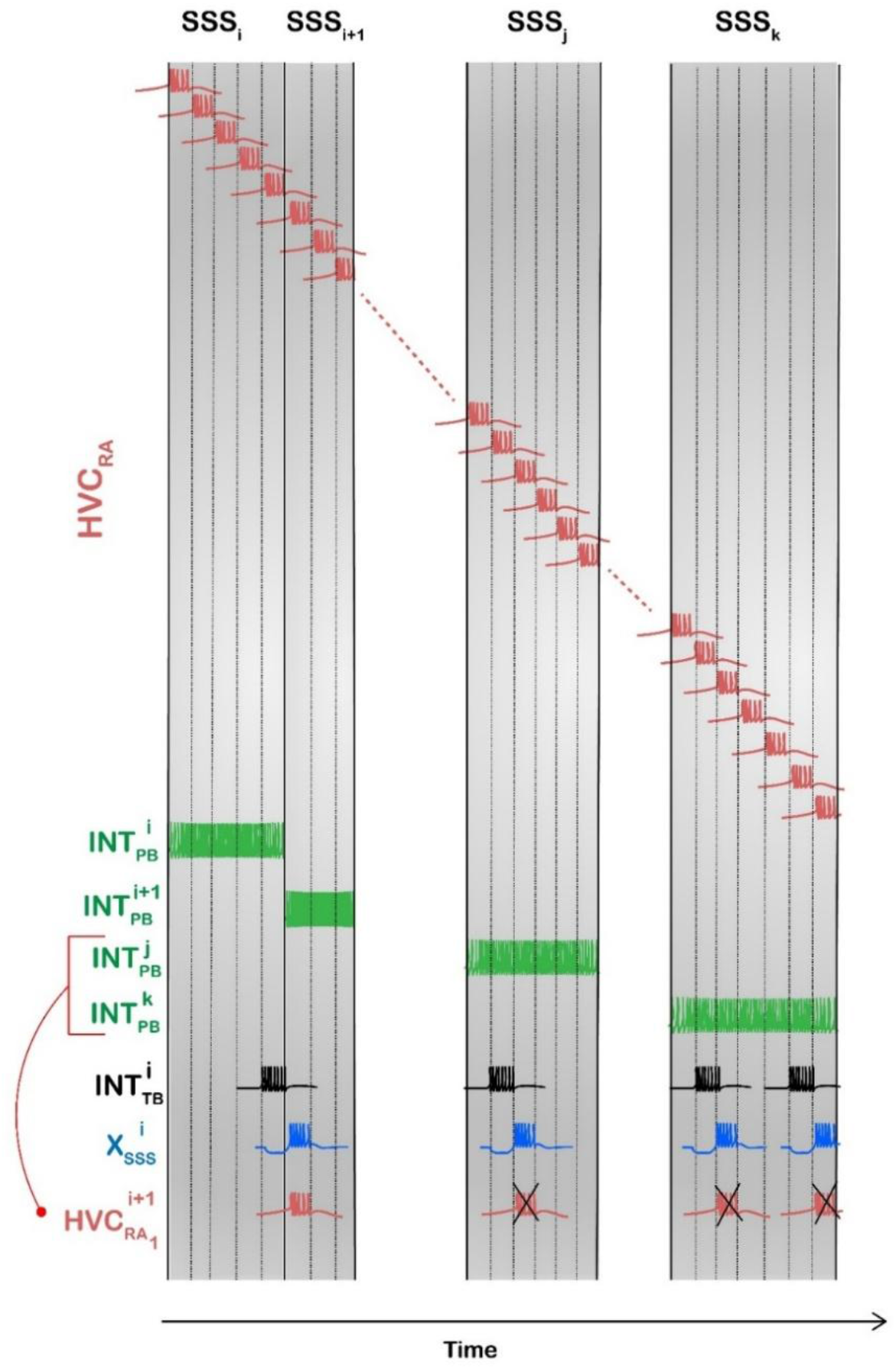
Functional logic for propagation and stability in the microcircuit chain. Representative membrane potential traces showing how microcircuit-specific phasic and tonic inhibition coordinates HVC_RA_ and HVC_X_ activity to advance the sequence while preventing spurious reactivation. Gray boxes depict four sub-syllabic segments (SSSs): two consecutive microcircuits (*SSS*_*i*_ and *SSS*_*i*+1_) and two later, nonconsecutive segments (*SSS*_*j*_ and *SSS*_*k*_). Top (**red**): within each SSS, a randomly sized HVC_RA_ ensemble (examples: 5, 3, 6, 7 cells) generates an ordered burst packet via feedforward AMPA chaining. **Green**: the MS phasic interneuron *INT*_*PB*_ integrates excitation from the local HVC_RA_ ensemble and produces a segment-spanning inhibitory epoch that defines the SSS temporal “window.” **Black:** at segment completion, the tonic interneuron 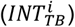 inhibits the MS X-projector (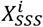, **blue**), producing a hyperpolarizing sag; release triggers a rebound burst that excites the first HVC_RA_ neuron of the next microcircuit (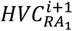, **red**) to initiate the next packet. Additional tonic bursts (driven by later-segment excitation) generate extra 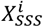 rebounds; phasic inhibition from the active segment 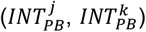 vetoes the resulting out-of-window activations of 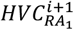 (X marks), preventing pathological restarts while preserving single-burst HVC_RA_ alongside multi-burst HVC_X_. Time runs left to right.

### HVC_RA_ packets define local segments; PB interneurons define the segment “window”

Within each microcircuit, sequential HVC_RA_ bursting is generated by directed AMPA coupling along the local HVC_RA_ chain (Figure 1), producing a brief, ordered packet aligned to that SSS. Because all HVC_RA_ neurons in a microcircuit excite its MS phasic interneuron (*INT*_*PB*_), *INT*_*PB*_ integrates the HVC_RA_ packet and produces a segment-spanning inhibitory burst. For example, in microcircuit *i*, the five HVC_RA_ neurons collectively drive 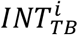, generating a “long” phasic burst whose onset coincides with the first spike of 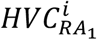 and whose offset tracks the last spike of 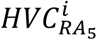, thereby defining the temporal window of *SSS*_*i*_ (Figure 2, green trace under *SSS*_*i*_). Functionally, this “windowing” role resembles feedforward inhibition in cortical circuits, where rapidly recruited interneurons impose a narrow time window for excitation to influence spiking and suppress late destabilizing drive^45,46^. In this sense, *INT*_*PB*_ structures time rather than simply suppressing activity.

### TB interneurons mark segment completion and recruit HVC_X_ rebound bursting to propagate forward

In contrast, the MS tonic interneuron (*INT*_*TB*_) is driven primarily by the terminal HVC_RA_ neuron of the local chain. In microcircuit *i*, 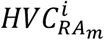 excites 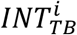,producing a burst near the end of *SSS*_*i*_. 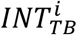 then inhibits the MS X-projecting neuron 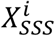, producing a pronounced hyperpolarizing “sag” in 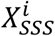 (Figure 2, blue). Release from this inhibition triggers a rebound burst in 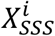, which excites the first HVC_RA_ neuron of the next microcircuit, 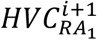, initiating the HVC_RA_ chain for *SSS*_*i*+1_. Thus, the backbone of propagation is: tonic inhibition → HVC_X_ rebound → next-segment HVC_RA_ recruitment, a conserved computational motif in which inhibition becomes a precisely timed trigger^47–49^.

This logic then repeats across microcircuits (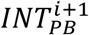 spans *SSS*_*i*+1_; 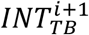 recruits rebound in 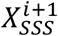; activity advances to microcircuit *i* + 2).

### Why two inhibitory modes are required: preventing restarts while allowing multi-burst HVC_X_

A key challenge is that the same relay that advances the sequence 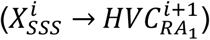 also creates vulnerability to spurious reactivation if 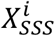 rebounds multiple times. In the model, 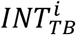 can be recruited not only by the terminal HVC_RA_ neuron in microcircuit *i*, but also by excitatory inputs from HVC_RA_ and HVC_X_ neurons in other microcircuits. In the example shown, 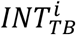 receives additional excitation from 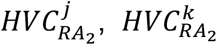 and 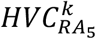, producing four tonic bursts across the motif (Figure 2, black trace), each inducing a corresponding inhibition-and-release event in 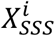 and thus multiple rebound bursts (Figure 2, blue). This naturally yields the experimentally observed multi-burst regime of HVC_X_ neurons (2-4 bursts per motif). Without additional control, each rebound burst would re-excite 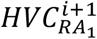 and re-trigger the HVC_RA_ chain in microcircuit *i* + 1, producing extra HVC_RA_ bursts that violate the hallmark single-burst regime and effectively mimic pathological “restart-like” dynamics. The model prevents this failure mode through the phasic inhibitory gate: for any segment *SSS*_*x*_, 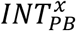 suppresses HVC_RA_ neurons attempting to fire outside their assigned window. Accordingly, the “extra” 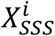 rebounds that occur during later segments (*SSS*_*j*_ and *SSS*_*k*_) are vetoed by 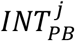 and 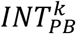, which silence 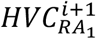 at those inappropriate times (Figure 2, red), preventing unwanted reactivation of microcircuit *i* + 1.

Together, these mechanisms assign complementary roles to inhibition: tonic inhibition recruits rebound bursting in HVC_X_ to provide a reliable forward-propagating trigger, whereas phasic inhibition enforces temporal exclusivity so that only the HVC_RA_ ensemble assigned to an SSS, bursts during that SSS. This division of labor reconciles multi-burst HVC_X_ with single-burst HVC_RA_ within a single propagating sequence and mirrors stabilization strategies in other sequence-generating circuits, where progression depends on release-from-inhibition while parallel inhibitory motifs prevent re-entrant dynamics^50,51^.

### Activity patterns of HVC_RA_ neurons

We first describe the population activity produced by the network (Figures 3–5). HVC_RA_ neurons burst with extreme sparsity, generating at most one burst per motif (typically 4-6 spikes). Across simulations, HVC_RA_ bursts lasted 7.94 ± 0.92 ms and contained 5.89 ± 0.74 spikes, reproducing the defining *in vivo* signature of HVC_RA_ during singing - brief, precisely time-locked bursting with a single event per motif time point^19,52^. Figure 3 visualizes this sparse code for 30 HVC_RA_ neurons aligned to a representative syllable segmented into five sub-syllabic segments (*SSS*_*i*_ - *SSS*_*i*+4_). Consistent with the random recruitment rule (3-10 HVC_RA_ neurons per microcircuit), the number of HVC_RA_ neurons assigned to each segment varies (here: 5, 3, 4, 12, and 6, respectively; vertical boundaries). Within each microcircuit, ordered HVC_RA_ bursting arises from feedforward excitation along the local HVC_RA_ chain (Figure 1), producing a compact packet confined to the segment window.

**Figure 3.**
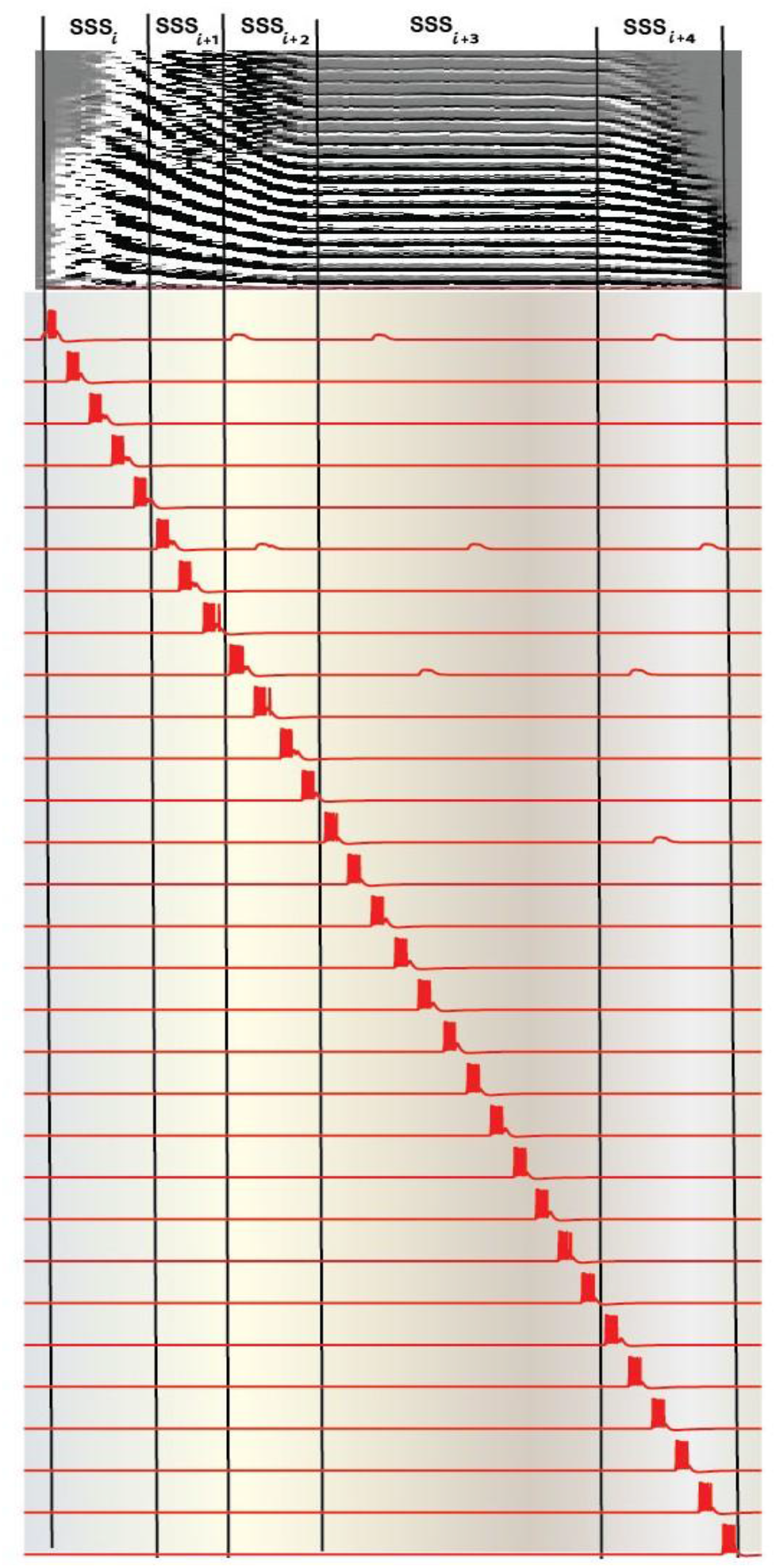
Ultra-sparse, segment-locked HVC_RA_ bursting in the model. **Top:** Example syllable waveform partitioned into five sub-syllabic segments (*SSS*_*i*_ - *SSS*_*i*+4_; boundaries indicated by vertical lines). **Bottom**: Membrane potential traces from 30 representative HVC_RA_ neurons aligned to the syllable. HVC_RA_ neurons emit at most one brief burst per motif (typically 4-6 spikes), with different neurons bursting at distinct times to form a sparse sequential packet. Differences in burst density across SSSs reflect random recruitment of 3-10 HVC_RA_ neurons per microcircuit. Subthreshold depolarizations (“bumps”), most prominent in chain-initiating HVC_RA_ neurons, arise from excitatory input driven by microcircuit-specific HVC_X_ rebound bursts; segment-specific phasic inhibition vetoes these inputs to prevent off-time HVC_RA_ reactivation (Figure 6).

**Figure 4.**
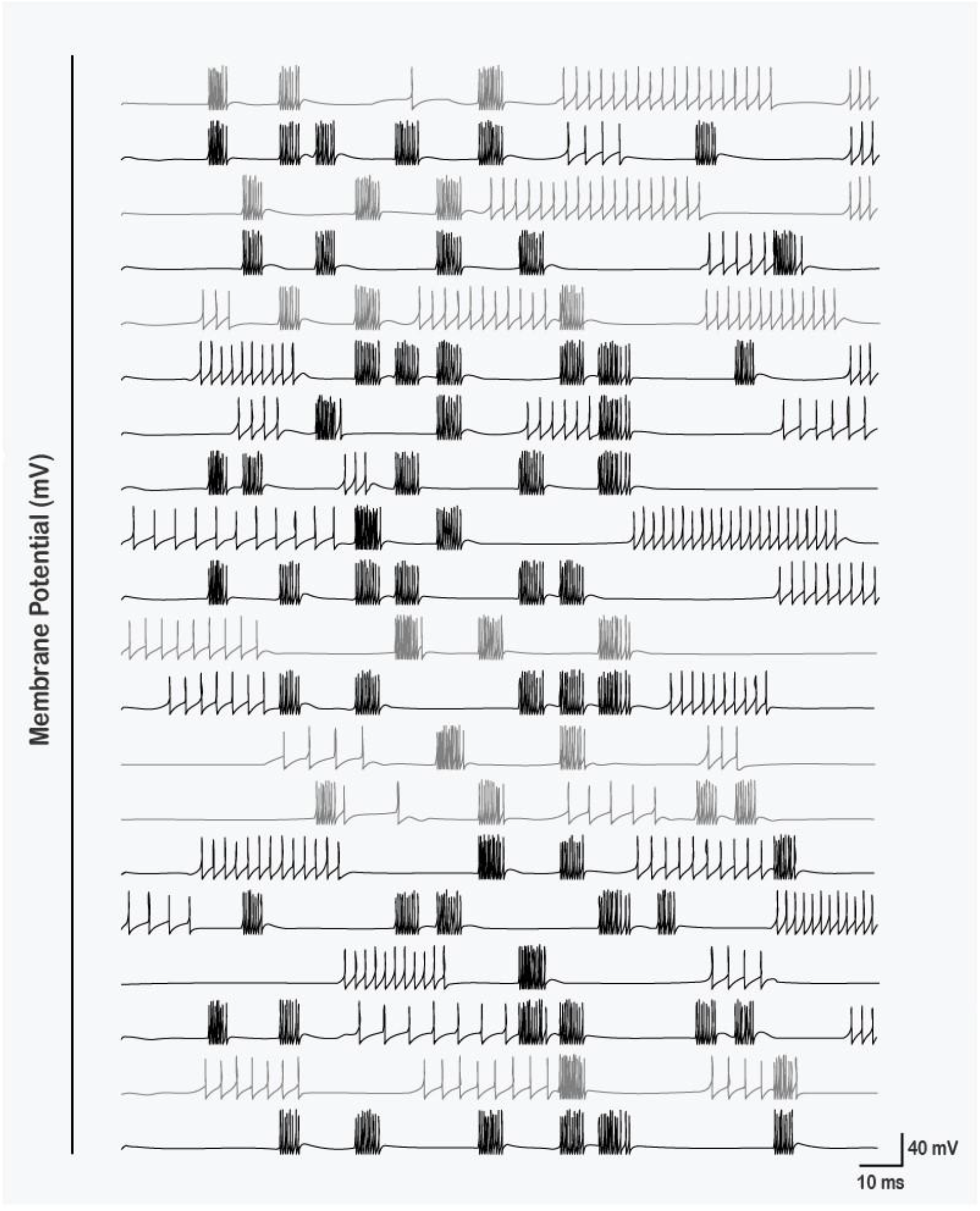
Motif-spanning bursting and irregular spiking in tonic HVC interneurons. Representative membrane potential traces from 20 tonic-bursting (TB) interneurons during a simulated motif, including microcircuit-specific cells (light gray) and non-microcircuit-specific cells (black). TB interneurons generate multiple bursts distributed across the motif, driven by convergent excitation from HVC_RA_ neurons (across microcircuits) and from both MS and NMS HVC_X_ neurons (Figure 2). Inter-burst irregular spiking arises from stochastic synaptic pulse trains (Methods), producing an *in vivo*-like inhibitory activity.

**Figure 5.**
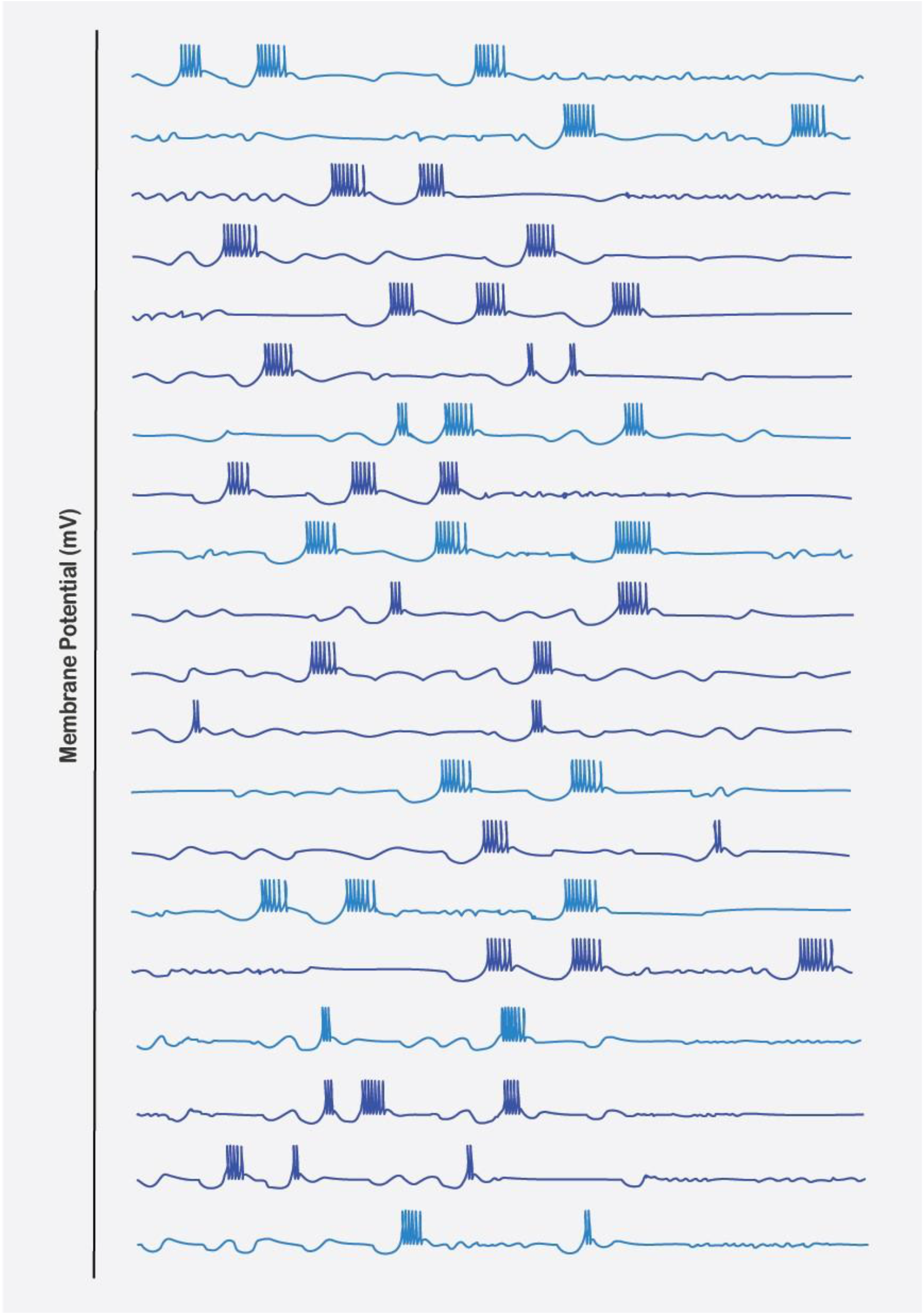
Rebound-driven multi-burst firing in HVC_X_ neurons. Representative membrane potential traces from 20 HVC_X_ neurons during a simulated motif, including microcircuit-specific (MS; light blue) and non-microcircuit-specific (NMS; dark blue) cells. HVC_X_ neurons generate 2-4 bursts with cell-to-cell variability in burst count and spikes per burst. In the model, bursts arise as post-inhibitory rebounds: MS HVC_X_ neurons are inhibited primarily by their assigned microcircuit-specific tonic-bursting interneuron, whereas NMS HVC_X_ neurons can be inhibited by multiple interneurons from the global pool. These rebound bursts provide a motif-scale timing signal that supports sequence propagation and aligns with experimentally observed multi-burst HVC_X_ firing.

Notably, the first HVC_RA_ neuron in each microcircuit exhibits small subthreshold “bumps,” reflecting excitatory drive from the corresponding microcircuit-specific HVC_X_ neuron. Because each *X*_*SSS*_ generates 2-4 rebound bursts (Figure 2), the first rebound burst providing the forward-propagating trigger to the next microcircuit and later bursts reflecting additional inhibition-release events from *INT*_*TB*_, these HVC_X_-driven depolarizations would, in principle, create unwanted HVC_RA_ reactivations and restart the local chain outside its assigned segment. In the model, these incipient events are vetoed by phasic inhibition: the segment-appropriate *INT*_*PB*_ suppresses HVC_RA_ spiking at off-time moments. For example, 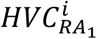 shows three bumps during *SSS*_*i*+2_ - *SSS*_*i*+4_ because upstream 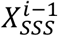 rebounds during those later segments, but spiking is silenced by 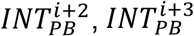 and 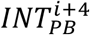. We dissect this inhibitory veto mechanism directly in Figure 6.

**Figure 6.**
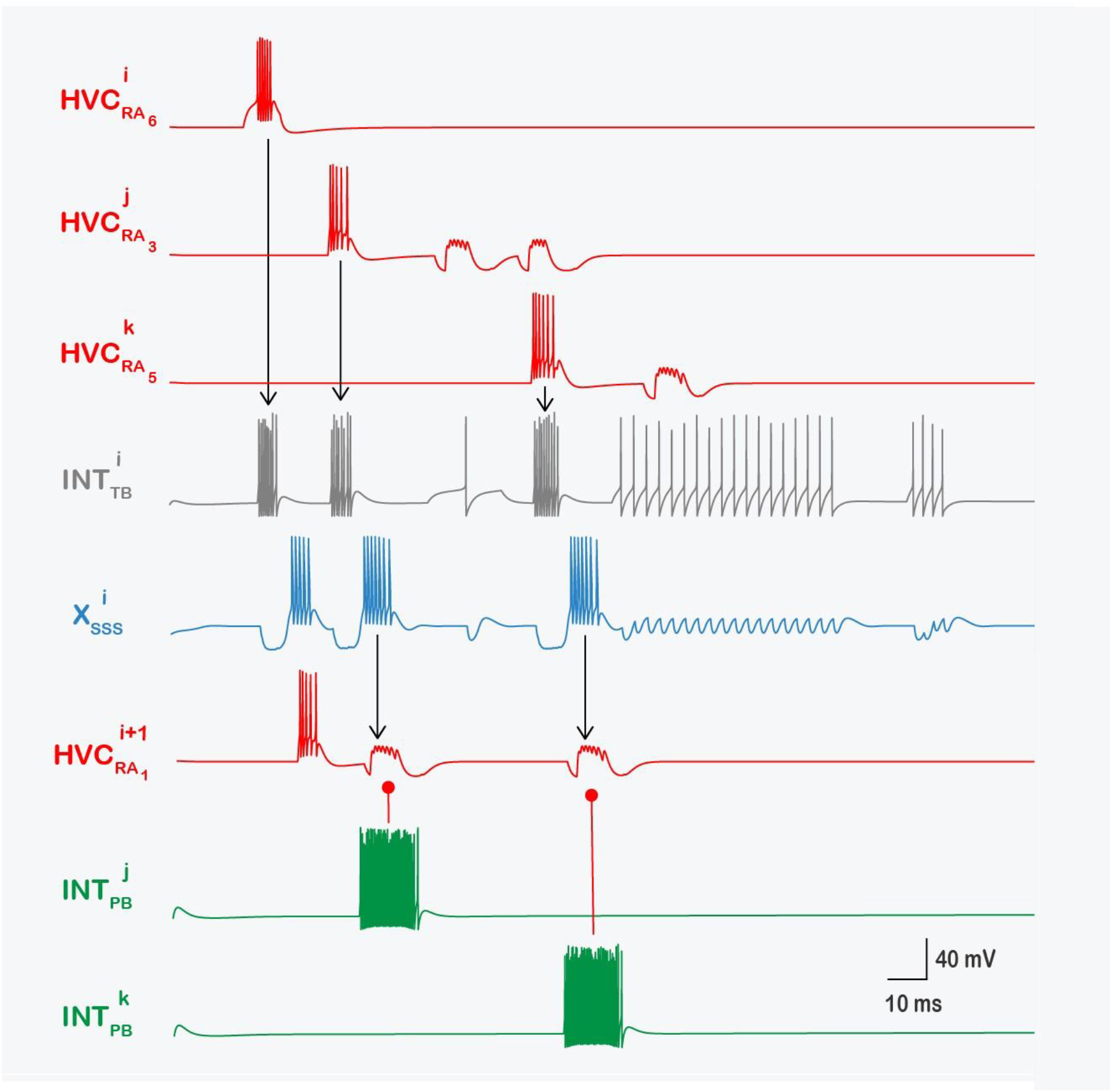
Microcircuit interplay: tonic inhibition recruits HVC_X_ rebound bursts for propagation, while phasic inhibition vetoes off-time HVC_RA_ reactivation. Representative membrane potential traces showing stabilizing interactions within and across microcircuits. Three HVC_RA_ neurons (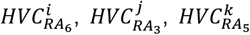; **red**) provide AMPA excitation to the microcircuit-specific tonic interneuron 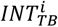 (**gray**), eliciting time-locked bursts that inhibit the microcircuit-specific HVC_X_ neuron 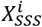 (**blue**) and generate post-inhibitory rebound bursts. The first rebound excites the next microcircuit’s chain-initiating neuron 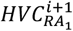 (red) to trigger correct propagation, whereas later rebounds would re-excite 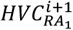 and induce non-biological extra HVC_RA_ bursts. Segment-specific phasic interneurons (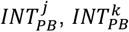; **green**) prevent these restarts via GABAergic inhibition onto 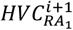, converting would-be spikes into subthreshold “bumps.”

### Determinants of HVC_RA_ burst strength, spike count, onset timing, and propagation robustness

Beyond enforcing sparsity, the model specifies how each HVC_RA_ burst - its onset latency, spike count, and duration - emerges from a constrained interaction between synaptic drive and intrinsic excitability, and shows that the same “control knobs” that tune burst shape also define the boundary between stable propagation and sequence failure. First, burst strength and onset latency are set by the effective AMPA conductance recruiting each HVC_RA_ neuron: either HVC_RA_ → HVC_RA_ within a microcircuit (for non-initial chain elements) or 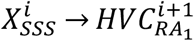 across the microcircuit boundary (for the chain-initiating neuron). In both cases, larger *g*_*AMPA*_ yields stronger bursts, more spikes, and shorter first-spike delays, consistent with synaptic control of recruitment timing in feedforward sequences. Second, spike onset and burst size are shaped intrinsically by the A-type K^+^ conductance (*g*_*A*_): increasing *g*_*A*_ delays recruitment and reduces spike count by lowering effective excitability. Third, burst termination is governed by the Ca^2+^-dependent K^+^ conductance (*g*_*SK*_): Ca^2+^entry through the L-type current (*I*_*CaL*_) accumulates during the burst, progressively recruiting *I*_*SK*_ until the outward current curtails spiking^7^. Together, these mechanisms give a biophysical account of why HVC_RA_ bursts remain brief and stereotyped while still being tunable by synaptic drive and intrinsic conductance state.

Crucially, proper sequence propagation is not guaranteed by architecture alone: it requires that these parameters remain within a bounded regime. Supplementary Figure 2 identifies three HVC_RA_-level failure modes that fracture propagation. First, within-microcircuit chaining fails if the AMPA conductance between successive HVC_RA_ neurons is sufficiently small that postsynaptic neurons cannot reach threshold. Second, propagation fails at microcircuit boundaries if 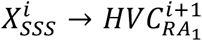 AMPA drive is too weak to recruit the next chain’s initiator (Supplementary Figure 2A). Third, even with intact synaptic connectivity, propagation can be eliminated when intrinsic “braking” currents push HVC_RA_ neurons outside a burst-permissive excitability regime: sufficiently large *g*_*A*_ or *g*_*SK*_ can abolish bursting altogether (Supplementary Figure 2B-C). Thus, robust HVC_RA_ propagation requires a minimum synaptic gain (both within-chain and X → HVC_RA_ coupling and a bounded intrinsic “braking” regime (particularly *g*_*A*_ and *g*_*SK*_).

### Activity patterns of HVC_INT_ neurons and tonic-interneuron failure modes that destabilize propagation

Tonic-bursting interneurons in the model exhibit motif-spanning, high-activity regimes that mirror the dense inhibitory backdrop observed in HVC during singing. Figure 4 shows representative membrane potential traces for 20 TB interneurons, including both microcircuit-specific cells (light black) and non-microcircuit-specific cells (dark black). Across the motif, TB interneurons generate multiple bursts, consistent with their role as broadly recruited inhibitory controllers rather than segment-local, single-event units. These bursts arise from convergent AMPA excitation driven by distributed projection-neuron activity, inputs from HVC_RA_ neurons (from any microcircuit) and from both microcircuit-specific (MS) and non-microcircuit-specific (NMS) HVC_X_ neurons (as in Figure 2), so TB interneurons effectively integrate motif-wide excitation and translate it into structured inhibitory epochs (quantified below; Figures 6-7). TB interneurons also show irregular spiking between (and outside) burst epochs due to stochastic synaptic pulse trains (Methods), capturing the synaptic “bombardment” regime characteristic of *in vivo* networks where ongoing fluctuations coexist with event-like structure^53–55^. In this circuit, burst epochs riding on an irregular inhibitory background provide robustness (ongoing inhibitory tone) without requiring interneurons to be silent outside assigned windows.

**Figure 7.**
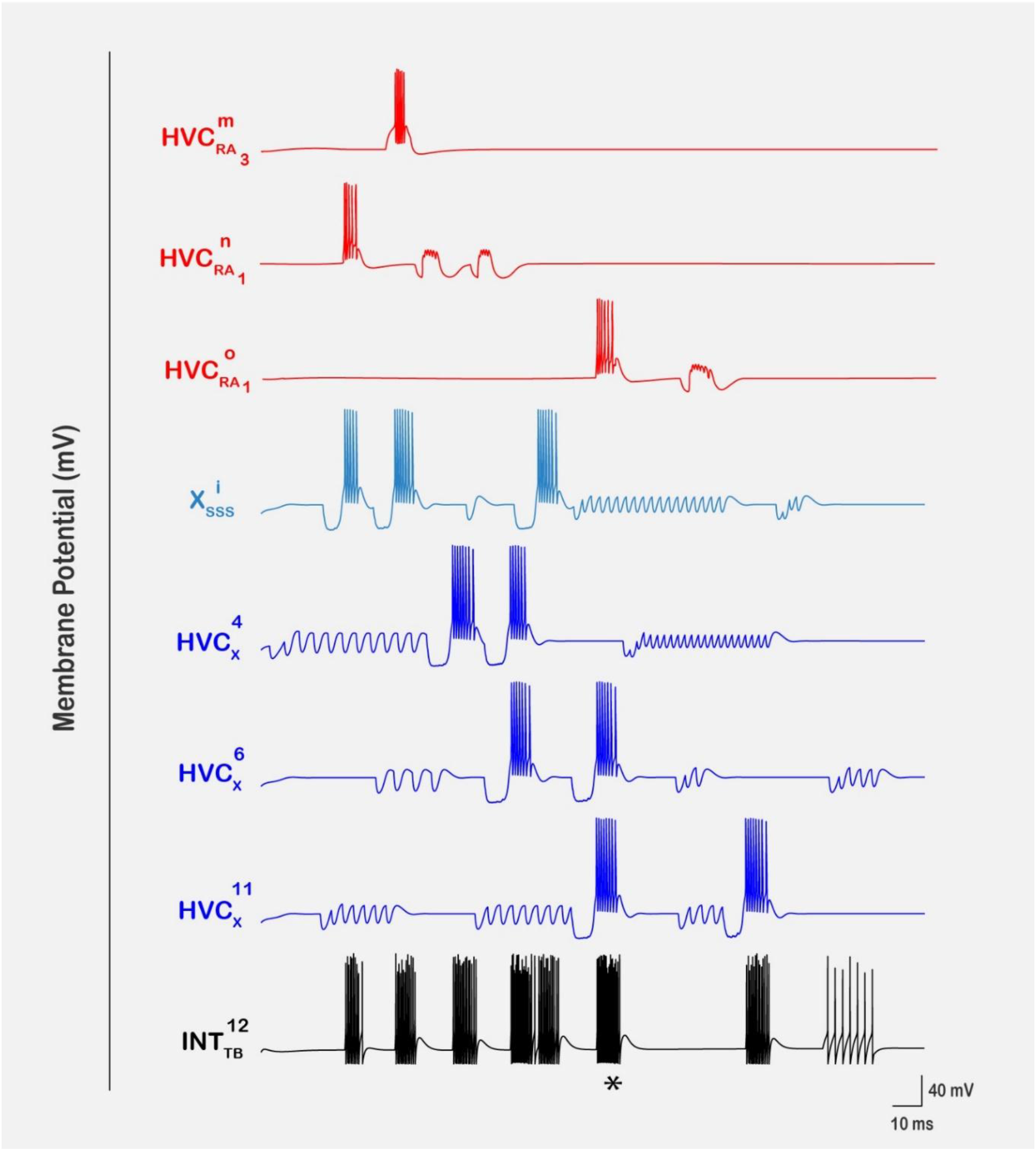
Distributed convergence from HVC_RA_ and HVC_X_ drives tonic bursting in HVC interneurons. Representative membrane potential traces illustrating how a tonic-bursting interneuron from the non-microcircuit-specific pool, 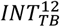 (black), is recruited by convergent excitation from both classes of projection neurons. 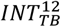receives AMPAergic inputs from three HVC_RA_ neurons (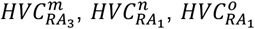; red) and from four HVC_X_ neurons, including one microcircuit-specific X-projector (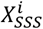; light blue) and three non-microcircuit-specific X-projectors (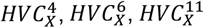; dark blue). Each presynaptic burst can recruit a time-locked burst in 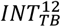, yielding motif-spanning multi-burst activity. The asterisk marks a particularly strong interneuron burst produced by near-coincident presynaptic bursting (here 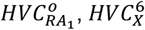 and 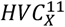), illustrating temporal summation of excitatory drive that increases interneuron firing frequency and spike count.

Critically, the model also shows that sequence propagation is highly sensitive to TB interneuron excitability, because TB output directly gates whether MS HVC_X_ neurons can escape inhibition to generate the rebound bursts that drive the HVC_X_ → HVC_RA_ relay, a pathway whose causal relevance is now strengthened by the new Trusel et al.^43^ findings. When excitation from projection neurons onto MS TB interneurons becomes too strong (≫ *g*_*AMPA*_), TB cells enter regimes of dense, near-continuous bursting/spiking, imposing prolonged inhibition onto MS X-projecters; this delays or halts rebound and thereby destabilizes (or stops) propagation. Similarly, perturbing intrinsic rebound-supporting conductances in MS TB interneurons can push them into runaway excitation: increasing *g*_*CaT*_ (3-fold) or *g*_*H*_ (5-fold) in 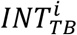 drives sustained TB activation that continuously inhibits 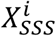, preventing rebound and blocking transmission to 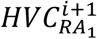, thereby halting the sequence (Supplementary Figure 3A-B). Mechanistically, these perturbations are particularly destabilizing because both channel types are powerful excitability controllers near rest: T-type Ca^2+^ channels open close to resting membrane potential and markedly enhance responsiveness when upregulated, while H-channels strongly shape baseline depolarizing drive and excitability (and can indirectly promote calcium entry by maintaining depolarized regimes). Thus, TB burst count, duration, and spikes-per-burst are not merely descriptive outputs, they are emergent consequences of synaptic drive and intrinsic excitability that must remain within a constrained regime to preserve inhibitory timing, enable HVC_X_ rebound, and maintain stable motif-scale propagation.

### Activity patterns of HVC_X_ neurons

HVC_X_ neurons generate 2-4 motif-locked bursts in the model (Figure 5; 20 cells shown), for both MS (light blue) and NMS (dark blue) populations, matching the core *in vivo* phenomenology of HVC_X_ during singing^19,22,26,27^. Mechanistically, HVC_X_ bursts are post-inhibitory rebound events: MS HVC_X_ neurons are inhibited primarily by their assigned MS tonic-bursting interneuron, whereas NMS HVC_X_ neurons can be inhibited by multiple interneurons, naturally producing heterogeneity in burst count and spikes-per-burst. This rebound mechanism provides a principled route for inhibition to create precisely timed excitation via deinactivation of T-type Ca^2+^ channels and recruitment of *I*_*h*_ ^47–49^. In HVC, this logic is especially consequential because rebound-generated HVC_X_ bursts can be converted directly into forward progression through the HVC_X_ → HVC_RA_ relay, now reinforced by recent connectivity mapping^43^: inhibition becomes a timing signal that launches the next segment rather than a merely destabilizing brake.

Because the projection from 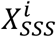 to 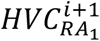 is the model’s critical relay for advancing from one sub-syllabic segment to the next (Figures 2 and 6), it is also a natural point of fragility. In the model, each rebound burst in 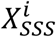 must be strong enough to drive 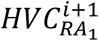 suprathreshold; otherwise, the next microcircuit is not recruited and propagation collapses at the boundary. Supplementary Figure 4 illustrates a synaptic route to relay failure: reducing the GABAergic conductance from the MS tonic interneuron to the MS X-projecter (*g*_*GABA*_, 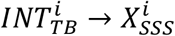) by 10-fold preserves the burst in 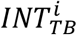 (still driven by the terminal RA neuron 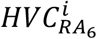) and produces a visible sag in 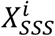, but release yields only a small rebound overshoot rather than a rebound burst. Consequently, 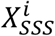 fails to recruit 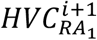, later microcircuits remain silent (e.g., 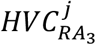 and 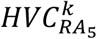), 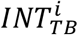 expresses only the single burst tied to microcircuit *i* (no later excitatory events arrive), and downstream phasic interneurons (e.g. 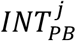 and 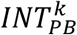) are silent because their local HVC_RA_ ensembles were never activated.

A second constraint is intrinsic: the relay is only as reliable as the ability of 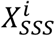 to generate a suprathreshold rebound burst. Supplementary Figure 5A shows that reducing the T-type Ca^2+^ conductance (*g*_*CaT*_) in 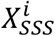 by 3-fold weakens/abolishes rebound bursting and collapses propagation with the same downstream signature as Supplementary Figure 4. Conversely, Supplementary Figure 5B shows that increasing the Ca^2+^-dependent K^+^ conductance (*g*_*SK*_) in 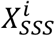 by 2-fold suppresses rebound excitability: the potentiated outward *I*_*SK*_ counteracts the inward rebound drive from *I*_*CaT*_ (with support from *I*_*h*_), preventing 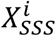 from reaching a burst-permissive regime and breaking the chain. Together, these perturbations define a bounded “rebound window” required for stable propagation, where inward rebound currents (dominated by *g*_*CaT*_, supported by *I*_*h*_) must overcome, but remain balanced by, activity-dependent outward stabilization (*g*_*SK*_).

Finally, the model identifies an opposite failure mode in which rebound gain becomes too large. Supplementary Figure 6 increases *g*_*H*_ in 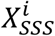 by 5-fold, pushing 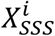 into runaway excitation: it fires nearly continuously, pausing only during inhibition from 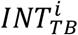. Through the 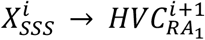 projection, this immediately drives 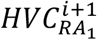 into continuous firing-pathological for HVC_RA_ - and the effect propagates through microcircuit *i* + 1, causing the remaining HVC_RA_ neurons to adopt similarly dense firing. This forces 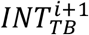 into high-frequency firing and strongly clamps 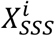 for most of the motif. Notably, 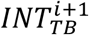 becomes silent only during the same brief window when 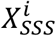 is silenced by 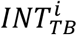; during that shared silent window 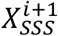 escapes inhibition and generates a single rebound burst that eventually activates 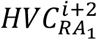, but at a markedly delayed time outside its intended SSS. Thus, excessive *I*_*h*_ need not simply “break” the chain, it can time-warp it, producing dense firing in HVC_RA_ (instead of ultra-sparse bursts), high-frequency firing in HVC_X_ (instead of 2-4 rebounds), delayed recruitment of downstream microcircuits (phase shifts), and in other parameter regimes complete sequence failure.

### Neuronal interplays within the microcircuit

We next dissect the tonic-phasic division of labor at the single-microcircuit level. In the example microcircuit *i* (Figure 6), the tonic interneuron 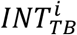 receives AMPA drive from the terminal HVC_RA_ neuron of microcircuit *i* (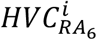 (top red; the segment-completion signal; Figures 2-3) and additional HVC_RA_ neurons activated later in the motif. As a result, 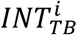 emits multiple time-locked bursts and shows irregular inter-burst spiking due to ongoing synaptic drive (Methods). Each 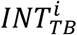 burst inhibits the MS HVC_X_ neuron 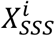, and release recruits a post-inhibitory rebound burst. The first rebound burst excites the next microcircuit’s chain-initiating HVC_RA_ neuron 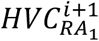 to launch the subsequent HVC_RA_ packet and advance the sequence. Later 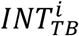-driven rebound bursts would, in principle, re-excite 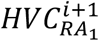; these off-time restarts are prevented by the phasic interneurons active in the current segment 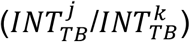, which veto HVC_RA_ spiking outside its assigned window (see Figure 2).

### Phasic-veto failure modes define a narrow operational regime

Weakening PB → HVC_RA_ inhibition undermines this stabilizing veto and allows ectopic HVC_RA_ bursts to escape. When *g*_*GABA*_ from a single phasic burster (e.g. 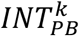) to its HVC_RA_ targets is reduced 5-fold (Supplementary Figure 7A), an off-time rebound-driven event that should be vetoed instead produces an ectopic burst in 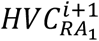, which then seeds downstream off-schedule activity through interneuron/X-projecter interactions. When PB → HVC_RA_ inhibition is reduced 5-fold globally across phasic bursters (Supplementary Figure 7B), the failure becomes circuit-wide: the network loses segment exclusivity, many HVC_RA_ neurons drift into inappropriate multi-burst/tonic-like regimes, and activity becomes re-entrant and non-song-like. Conversely, the model shows that PB neurons must also avoid excessive intrinsic excitability: increasing *g*_*CaT*_ 4-fold or *g*_*H*_ 5-fold in a representative phasic burster 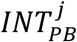 (Supplementary Figure 8A-B) converts a brief, segment-locked veto into a persistently active inhibitor, imposing an effectively continuous clamp on its postsynaptic HVC_RA_ targets and locally abolishing legitimate segment bursts. Thus, stable propagation requires PB interneurons to remain within a constrained regime: strong enough to veto off-time reactivation, but not so excitable that they extinguish the sparse, segment-locked HVC_RA_ code they are meant to protect.

### Tonicity in the bursting activity of HVC_INT_ neurons

Figure 7 shows how tonic-bursting interneurons (*INT*_*TB*_) acquire motif-spanning “tonicity” through distributed convergent excitation. We highlight one representative interneuron from the non-microcircuit-specific (NMS) pool, 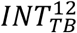, alongside the activity of all presynaptic projection neurons that contact it. In this example, 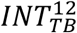 receives random AMPAergic inputs from three HVC_RA_ neurons 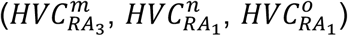 and four HVC_X_ neurons: one microcircuit-specific projector 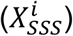 and three NMS X-projectors 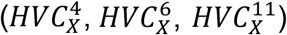. This convergence recruits multiple interneuron bursts distributed across the motif.

More generally, in the model the number of *INT*_*TB*_ bursts are set by the burst “opportunities” provided by presynaptic partners: each presynaptic HVC_RA_ or HVC_X_ burst can recruit a time-locked interneuron burst. The strength of each interneuron burst (spike count/firing frequency) is governed primarily by the effective AMPA conductance from projection neurons onto *INT*_*TB*_, such that stronger coupling produces more powerful bursts. The model further predicts a nonlinear summation regime: temporally overlapping presynaptic bursts drive disproportionately strong interneuron responses. This is illustrated by the asterisk in Figure 7, where coincident bursts in 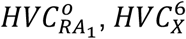 and 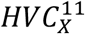 yield a markedly stronger, denser burst in 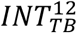. In the next section, we show that intrinsic conductances controlling interneuron excitability can further reshape recruitment and, when perturbed, destabilize sequence propagation.

### Rebound bursting in HVC_X_ neurons

HVC_X_ bursting in the model is governed by a tight coupling between synaptic and intrinsic mechanisms, which together set both how many rebound bursts occur and how strong each burst becomes. Synaptically, rebound amplitude scales with the depth, duration, and synchrony of inhibition: larger GABA_A_ conductances, greater temporal overlap among inhibitory interneuron bursts, and longer/denser inhibitory epochs all potentiate rebound, converting release from inhibition into either a weak depolarization or a full high-frequency burst. Thus, inhibition does not merely silence HVC_X_, it sets the initial conditions for the rebound event. Intrinsically, rebound is amplified primarily by T-type Ca^2+^ current and *I*_*h*_, which are progressively recruited by hyperpolarization (via deinactivation/activation) and then provide inward drive at release. Burst spike count is constrained by an opposing stabilizer, *g*_*SK*_, which dampens excitability and limits spikes-per-burst. Together, HVC_X_ bursts emerge from a push-pull interaction: inhibition-recruited inward currents (*I*_*CaT*_, *I*_*h*_) versus activity-dependent outward stabilization (*I*_*SK*_).

Figure 8A illustrates these dynamics for an exemplar NMS X-projecting neuron, 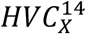, inhibited by three tonic interneurons 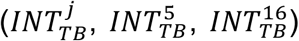. Periods of hyperpolarizing “sags” coincide with interneuron bursting and are summarized by bars (orange: strong/high-frequency inhibition; pink: weaker/sparser inhibition). Because inhibitory events often arrive in successive or overlapping bursts, 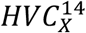 frequently remains trapped below threshold; brief gaps between inhibitory epochs yield only small rebound depolarizations, implying the release window is too short for *I*_*CaT*_ and *I*_*h*_ to fully engage. By contrast, two longer release intervals follow sufficiently strong/overlapping inhibition and allow *I*_*CaT*_ + *I*_*h*_ to reach burst-permissive states, producing robust rebound bursts, with *g*_*SK*_ shaping burst termination and spike count. Functionally, this inhibition-to-rebound transform is attractive for sequence generation because it allows inhibitory networks to act as *event setters*: the circuit can encode “when” a burst should occur by controlling the depth, duration, and synchrony of inhibition, rather than relying solely on excitatory ramping.

**Figure 8.**
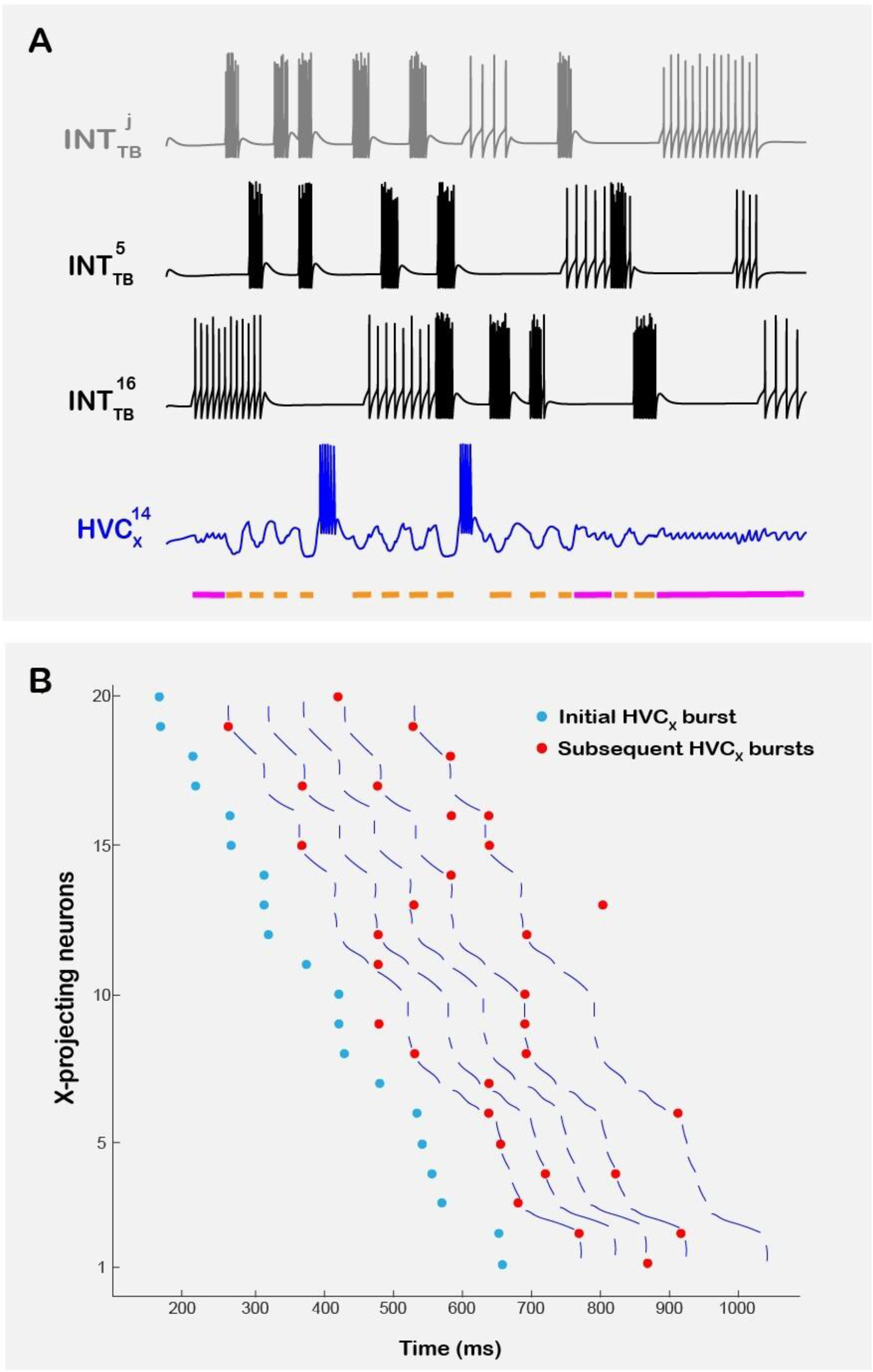
Inhibition-sculpted rebound bursting generates motif-locked HVC_X_ activity and parallel burst manifolds. (**A**). Example non-microcircuit-specific X-projecting neuron (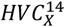, blue) illustrating post-inhibitory rebound bursting driven by convergent inhibition from three tonic-bursting interneurons (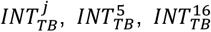; gray/black). Periods of interneuron bursting produce hyperpolarizing sags in 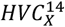 and suppress spiking; release from inhibition yields rebound depolarizations and occasional rebound bursts. Colored bars beneath the 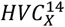 trace indicate epochs of inhibition: **orange** denotes strong inhibition associated with high-frequency interneuron bursting, whereas **pink** denotes weaker inhibition associated with sparse interneuron firing. Only sufficiently strong/prolonged inhibition followed by a sufficiently long silent interval permits a full rebound burst, consistent with rebound recruitment by intrinsic currents (e.g., *I*_*CaT*_, *I*_*h*_). (**B**) “Neuron cascade” representation of bursting in microcircuit-specific HVC_X_ neurons (*X*_*SSS*_) aligned to the canonical motif time base. Each row corresponds to a single neuron; neurons are ordered by the time of their first burst. Each marker denotes the time of the first spike in a burst: **blue** circles indicate the initial burst for each neuron and **red** circles indicate subsequent bursts. Initial bursts form a quasi-linear manifold across neurons, and later bursts form additional quasi-linear manifolds that run approximately parallel, recapitulating the “parallel lines” of HVC_X_ bursting reported experimentally (Fetterman & Margoliash, 2023). Lines illustrate the trajectory of bursts across successive events for each neuron, highlighting motif-spanning organization of multiple HVC_X_ burst sequences.

### Parallel “burst manifolds” in HVC_X_ emerge naturally from the microcircuit chain

Recent reanalysis indicates that HVC_X_ bursts can organize into multiple motif-spanning sequences (“parallel lines”), suggesting structured time coding in HVC_X_ alongside HVC_RA_’s ultra-sparse code^27^. To compare directly, we constructed the same “neuron cascade” (Figure 8B): bursts from each HVC_X_ neuron are plotted on one row; neurons are sorted by first-burst time; first bursts are blue and later bursts red; each marker indicates the first spike time of that burst.

Applied to microcircuit-specific HVC_X_ neurons, the model reproduces the key signature: first bursts form a quasi-linear manifold spanning the motif, and later bursts form additional quasi-linear manifolds approximately parallel to the first. This structure is emergent, not imposed by alignment rules, and matches the qualitative organization reported *in vivo*^27^. We focus on microcircuit-specific HVC_X_ neurons because the architecture supplies a mechanistic explanation: their rebound timing is organized by repeated, stereotyped inhibitory scheduling within the microcircuit chain. NMS HVC_X_ neurons can generate bursts that sometimes fall near these manifolds, but because their inhibitory drive is random (PB/TB inputs from the global pool), the model does not assign a unique functional origin to that alignment.

Supplementary Figure 9 provides a schematic account of how parallel manifolds arise. In brief, offsets between first bursts of successive *X*_*SSS*_ neurons reflect the propagation time through each microcircuit (set by SSS duration and HVC_RA_ chain length), while later bursts are produced by additional *INT*_*TB*_ - driven inhibitory events that recur at reproducible motif phases. This combination - segment timing plus repeated inhibitory scheduling - naturally generates parallel, shifted “passes” of rebound bursting and yields testable predictions for how manipulations that stretch/compress SSS duration or alter *INT*_*TB*_ recruitment should reshape manifold spacing.

## Discussion

We developed a biophysically grounded network model of zebra finch HVC that resolves a long-standing mechanistic tension: how ultra-sparse, single-burst HVC_RA_ activity can coexist with multi-burst HVC_X_ firing while still propagating a motif-precise sequence. The key solution is to treat inhibition as an active timing resource. Tonic interneuron bursts schedule post-inhibitory rebound in HVC_X_, and the resulting HVC_X_ → HVC_RA_ excitation provides a forward trigger at each segment boundary, while phasic inhibition defines segment-specific windows that veto off-time reactivation and preserve the “one-time-only” HVC_RA_ code.

This framework is strengthened by recent synaptic mapping in HVC indicating robust monosynaptic HVC_X_ → HVC_RA_ excitation and demonstrating that brief intra-HVC excitation can evoke restart-like motifs^56^. In our model, the same relay that enables reliable forward progression is also the natural locus of fragility: if rebound output is weakened (e.g., reduced inhibition onto *X*_*SSS*_, reduced *g*_*CaT*_, or elevated *g*_*SK*_), propagation fails; if rebound gain is pushed too high (e.g., excessive *I*_*h*_), the circuit transitions toward pathological dense firing and timing drift. Thus, stability emerges from a bounded rebound regime, not wiring alone.

Conceptually, the model unifies motifs that recur across timing circuits, release-from-inhibition generating precisely timed events and inhibitory windowing enforcing exclusivity, but ties them to identified HVC cell types and conductances. This cell-type grounding also provides an interpretable account of higher-order population structure: repeated inhibitory scheduling produces multiple motif-spanning “passes” of HVC_X_ rebound bursting, yielding parallel burst manifolds reminiscent of recent reanalysis of HVC_X_ activity^27^.

The model makes concrete, testable predictions. Increasing AMPA strength along HVC_RA_ chains or across the HVC_X_ → HVC_RA_ relay should advance recruitment and increase HVC_RA_ spike count, whereas elevating *g*_*A*_ should delay or veto HVC_RA_ recruitment within a segment. Manipulations that enhance *I*_*CaT*_/*I*_*h*_ in HVC_X_ neurons should increase rebound probability and burst strength, while increasing *g*_*SK*_ (or reducing Ca^2+^ entry) should truncate bursts and can break propagation at segment boundaries. Finally, weakening PB → HVC_RA_ inhibition should selectively permit off-window HVC_RA_ bursts and induce localized timing errors, whereas excessive PB excitability should over-clamp HVC_RA_ targets and abolish legitimate segment activity.

Several extensions remain. The present work focuses on mechanism rather than learning; incorporating plasticity rules that tune synaptic gains and intrinsic conductances could connect the rebound, veto architecture to how birds acquire and adapt song. Future work should also integrate interactions with upstream/downstream structures (e.g., NIf/Uva inputs and the basal ganglia loop), and explore how heterogeneity across interneuron subtypes and non-microcircuit-specific populations modulates robustness. More broadly, the model illustrates how conserved cellular mechanisms - post-inhibitory rebound and precisely timed inhibition - can be assembled into modular microcircuits that generate learned sequential behavior, offering testable predictions for how HVC implements millisecond-precise control in birdsong and, potentially, how analogous mechanisms support sequential computation in other vertebrate motor and cognitive systems.

## Notes

### Competing Interest Statement

The authors have declared no competing interest.

### Summary of Updates

There was an error in the author name as it appears in the PDF manuscript file: the name of my PhD student was entered incorrectly in the document itself. The author name is listed correctly however on the bioRxiv website, so the issue appears to be limited to the PDF file only. In short, the first author's name is Zeina Bou Diab and not Zeina Merabi (I happen to have two PhD students in my lab with the same first name, which made the confusion). To correct this, I am attaching the manuscript file with the author name entered properly. Nothing changed except the first author's last name.

### Funding

- University Research Board

## References

1. Skaggs, W.E., McNaughton, B.L., Wilson, M.A., and Barnes, C.A. (1996). Theta phase precession in hippocampal neuronal populations and the compression of temporal sequences. Hippocampus 6, 149–172. 10.1002/(SICI)1098-1063(1996)6:2%253C149::AID-HIPO6%253E3.0.CO;2-K.

2. Pastalkova, E., Itskov, V., Amarasingham, A., and Buzsáki, G. (2008). Internally generated cell assembly sequences in the rat hippocampus. Science 321, 1322–1327. 10.1126/science.1159775.

3. MacDonald, C.J., Lepage, K.Q., Eden, U.T., and Eichenbaum, H. (2011). Hippocampal “time cells” bridge the gap in memory for discontiguous events. Neuron 71, 737–749. 10.1016/j.neuron.2011.07.012.

4. Harvey, C.D., Coen, P., and Tank, D.W. (2012). Choice-specific sequences in parietal cortex during a virtual-navigation decision task. Nature 484, 62–68. 10.1038/nature10918.

5. Buzsáki, G., and Tingley, D. (2018). Space and Time: The Hippocampus as a Sequence Generator. Trends Cogn Sci 22, 853–869. 10.1016/j.tics.2018.07.006.

6. Peters, A.J., Chen, S.X., and Komiyama, T. (2014). Emergence of reproducible spatiotemporal activity during motor learning. Nature 510, 263–267. 10.1038/nature13235.

7. Bou Diab, Z., Chammas, M., and Daou, A. (2025). Biophysical network modeling of temporal and stereotyped sequence propagation of neural activity in the premotor nucleus HVC. Elife 14. 10.7554/eLife.105526.

8. Reiner, A., Laverghetta, A.V., Meade, C.A., Cuthbertson, S.L., and Bottjer, S.W. (2004). An immunohistochemical and pathway tracing study of the striatopallidal organization of area X in the male zebra finch. J Comp Neurol 469, 239–261. 10.1002/cne.11012.

9. Fee, M.S., and Scharff, C. (2010). The songbird as a model for the generation and learning of complex sequential behaviors. ILAR J 51, 362–377. 10.1093/ilar.51.4.362.

10. Vu, E.T., Mazurek, M.E., and Kuo, Y.C. (1994). Identification of a forebrain motor programming network for the learned song of zebra finches. J Neurosci 14, 6924–6934. 10.1523/JNEUROSCI.14-11-06924.1994.

11. Yu, A.C., and Margoliash, D. (1996). Temporal hierarchical control of singing in birds. Science 273, 1871–1875. 10.1126/science.273.5283.1871.

12. Fee, M.S., Kozhevnikov, A.A., and Hahnloser, R.H.R. (2004). Neural mechanisms of vocal sequence generation in the songbird. Ann N Y Acad Sci 1016, 153–170. 10.1196/annals.1298.022.

13. Long, M.A., and Fee, M.S. (2008). Using temperature to analyse temporal dynamics in the songbird motor pathway. Nature 456, 189–194. 10.1038/nature07448.

14. Mooney, R. (2009). Neurobiology of song learning. Curr Opin Neurobiol 19, 654–660. 10.1016/j.conb.2009.10.004.

15. Fee, M.S., and Goldberg, J.H. (2011). A hypothesis for basal ganglia-dependent reinforcement learning in the songbird. Neuroscience 198, 152–170. 10.1016/j.neuroscience.2011.09.069.

16. Vicario, D.S. (1991). Neural mechanisms of vocal production in songbirds. Curr Opin Neurobiol 1, 595–600. 10.1016/s0959-4388(05)80034-0.

17. Bolhuis, J.J., Okanoya, K., and Scharff, C. (2010). Twitter evolution: converging mechanisms in birdsong and human speech. Nat Rev Neurosci 11, 747–759. 10.1038/nrn2931.

18. Hahnloser, R.H.R., Kozhevnikov, A.A., and Fee, M.S. (2002). An ultra-sparse code underlies the generation of neural sequences in a songbird. Nature 419, 65–70. 10.1038/nature00974.

19. Kozhevnikov, A.A., and Fee, M.S. (2007). Singing-related activity of identified HVC neurons in the zebra finch. J Neurophysiol 97, 4271–4283. 10.1152/jn.00952.2006.

20. Fujimoto, H., Hasegawa, T., and Watanabe, D. (2011). Neural coding of syntactic structure in learned vocalizations in the songbird. J Neurosci 31, 10023–10033. 10.1523/JNEUROSCI.1606-11.2011.

21. Long, M.A., Jin, D.Z., and Fee, M.S. (2010). Support for a synaptic chain model of neuronal sequence generation. Nature 468, 394–399. 10.1038/nature09514.

22. Amador, A., Perl, Y.S., Mindlin, G.B., and Margoliash, D. (2013). Elemental gesture dynamics are encoded by song premotor cortical neurons. Nature 495, 59–64. 10.1038/nature11967.

23. Cannon, J., Kopell, N., Gardner, T., and Markowitz, J. (2015). Neural Sequence Generation Using Spatiotemporal Patterns of Inhibition. PLoS Comput Biol 11, e1004581. 10.1371/journal.pcbi.1004581.

24. Kosche, G., Vallentin, D., and Long, M.A. (2015). Interplay of inhibition and excitation shapes a premotor neural sequence. J Neurosci 35, 1217–1227. 10.1523/JNEUROSCI.4346-14.2015.

25. Markowitz, J.E., Liberti, W.A. 3rd, Guitchounts, G., Velho, T., Lois, C., and Gardner, T.J. (2015). Mesoscopic patterns of neural activity support songbird cortical sequences. PLoS Biol 13, e1002158. 10.1371/journal.pbio.1002158.

26. Lynch, G.F., Okubo, T.S., Hanuschkin, A., Hahnloser, R.H.R., and Fee, M.S. (2016). Rhythmic Continuous-Time Coding in the Songbird Analog of Vocal Motor Cortex. Neuron 90, 877–892. 10.1016/j.neuron.2016.04.021.

27. Fetterman, G.C., and Margoliash, D. (2023). Rhythmically bursting songbird vocomotor neurons are organized into multiple sequences, suggesting a network/intrinsic properties model encoding song and error, not time. Preprint, https://doi.org/10.1101/2023.01.23.525213 10.1101/2023.01.23.525213.

28. Dutar, P., Vu, H.M., and Perkel, D.J. (1998). Multiple cell types distinguished by physiological, pharmacological, and anatomic properties in nucleus HVc of the adult zebra finch. J Neurophysiol 80, 1828–1838. 10.1152/jn.1998.80.4.1828.

29. Kubota, M., and Taniguchi, I. (1998). Electrophysiological characteristics of classes of neuron in the HVc of the zebra finch. J Neurophysiol 80, 914–923. 10.1152/jn.1998.80.2.914.

30. Mooney, R. (2000). Different subthreshold mechanisms underlie song selectivity in identified HVc neurons of the zebra finch. J Neurosci 20, 5420–5436. 10.1523/JNEUROSCI.20-14-05420.2000.

31. Wild, J.M., Williams, M.N., Howie, G.J., and Mooney, R. (2005). Calcium-binding proteins define interneurons in HVC of the zebra finch (Taeniopygia guttata). J Comp Neurol 483, 76–90. 10.1002/cne.20403.

32. Mooney, R., and Prather, J.F. (2005). The HVC microcircuit: the synaptic basis for interactions between song motor and vocal plasticity pathways. J Neurosci 25, 1952–1964. 10.1523/JNEUROSCI.3726-04.2005.

33. Shea, S.D., Koch, H., Baleckaitis, D., Ramirez, J.-M., and Margoliash, D. (2010). Neuron-specific cholinergic modulation of a forebrain song control nucleus. J Neurophysiol 103, 733–745. 10.1152/jn.00803.2009.

34. Daou, A., Ross, M.T., Johnson, F., Hyson, R.L., and Bertram, R. (2013). Electrophysiological characterization and computational models of HVC neurons in the zebra finch. J Neurophysiol 110, 1227–1245. 10.1152/jn.00162.2013.

35. Drew, P.J., and Abbott, L.F. (2003). Model of song selectivity and sequence generation in area HVc of the songbird. J Neurophysiol 89, 2697–2706. 10.1152/jn.00801.2002.

36. Jin, D.Z. (2009). Generating variable birdsong syllable sequences with branching chain networks in avian premotor nucleus HVC. Phys Rev E Stat Nonlin Soft Matter Phys 80, 051902. 10.1103/PhysRevE.80.051902.

37. Gibb, L., Gentner, T.Q., and Abarbanel, H.D.I. (2009). Brain stem feedback in a computational model of birdsong sequencing. J Neurophysiol 102, 1763–1778. 10.1152/jn.91154.2008.

38. Gibb, L., Gentner, T.Q., and Abarbanel, H.D.I. (2009). Inhibition and recurrent excitation in a computational model of sparse bursting in song nucleus HVC. J Neurophysiol 102, 1748–1762. 10.1152/jn.00670.2007.

39. Hamaguchi, K., Tanaka, M., and Mooney, R. (2016). A Distributed Recurrent Network Contributes to Temporally Precise Vocalizations. Neuron 91, 680–693. 10.1016/j.neuron.2016.06.019.

40. Galvis, D., Wu, W., Hyson, R.L., Johnson, F., and Bertram, R. (2018). Interhemispheric dominance switching in a neural network model for birdsong. J Neurophysiol 120, 1186–1197. 10.1152/jn.00153.2018.

41. Egger, R., Tupikov, Y., Elmaleh, M., Katlowitz, K.A., Benezra, S.E., Picardo, M.A., Moll, F., Kornfeld, J., Jin, D.Z., and Long, M.A. (2020). Local Axonal Conduction Shapes the Spatiotemporal Properties of Neural Sequences. Cell 183, 537-548.e12. 10.1016/j.cell.2020.09.019.

42. Elmaleh, M., Kranz, D., Asensio, A.C., Moll, F.W., and Long, M.A. (2021). Sleep replay reveals premotor circuit structure for a skilled behavior. Neuron 109, 3851-3861.e4. 10.1016/j.neuron.2021.09.021.

43. Trusel, M., Zuo, J., Alam, D.H., Marks, E.S., Koch, T.M.I., Cao, J., Pancholi, H., Zhao, Z., Cooper, B.G., Zhang, W.-H., et al. (2026). Holistic motor control of zebra finch song syllable sequences. Nature. 10.1038/s41586-025-10069-z.

44. Kornfeld, J., Benezra, S.E., Narayanan, R.T., Svara, F., Egger, R., Oberlaender, M., Denk, W., and Long, M.A. (2017). EM connectomics reveals axonal target variation in a sequence-generating network. Elife 6, e24364. 10.7554/eLife.24364.

45. Pouille, F., and Scanziani, M. (2001). Enforcement of temporal fidelity in pyramidal cells by somatic feed-forward inhibition. Science 293, 1159–1163. 10.1126/science.1060342.

46. Isaacson, J.S., and Scanziani, M. (2011). How inhibition shapes cortical activity. Neuron 72, 231–243. 10.1016/j.neuron.2011.09.027.

47. Llinás, R., and Jahnsen, H. (1982). Electrophysiology of mammalian thalamic neurones in vitro. Nature 297, 406–408. 10.1038/297406a0.

48. Huguenard, J.R. (1996). Low-threshold calcium currents in central nervous system neurons. Annu Rev Physiol 58, 329–348. 10.1146/annurev.ph.58.030196.001553.

49. McCormick, D.A., and Bal, T. (1997). Sleep and arousal: thalamocortical mechanisms. Annu Rev Neurosci 20, 185–215. 10.1146/annurev.neuro.20.1.185.

50. Marder, E., and Bucher, D. (2001). Central pattern generators and the control of rhythmic movements. Curr Biol 11, R986–996. 10.1016/s0960-9822(01)00581-4.

51. Grillner, S. (2006). Neuronal networks in motion from ion channels to behaviour. An R Acad Nac Med (Madr) 123, 297–298.

52. Hahnloser, R.H.R., Kozhevnikov, A.A., and Fee, M.S. (2002). An ultra-sparse code underlies the generation of neural sequences in a songbird. Nature 419, 65–70. 10.1038/nature00974.

53. Softky, W.R., and Koch, C. (1993). The highly irregular firing of cortical cells is inconsistent with temporal integration of random EPSPs. J Neurosci 13, 334–350. 10.1523/JNEUROSCI.13-01-00334.1993.

54. Shadlen, M.N., and Newsome, W.T. (1998). The variable discharge of cortical neurons: implications for connectivity, computation, and information coding. J Neurosci 18, 3870–3896. 10.1523/JNEUROSCI.18-10-03870.1998.

55. Destexhe, A., Rudolph, M., and Paré, D. (2003). The high-conductance state of neocortical neurons in vivo. Nat Rev Neurosci 4, 739–751. 10.1038/nrn1198.

56. Trusel, M., Zhao, Z., Alam, D.H., Marks, E.S., Ikeda, M.Z., and Roberts, T.F. (2025). Synaptic connectivity of sensorimotor circuits for vocal imitation in the songbird. Elife 14. 10.7554/eLife.104609.

