## Supplementary material for "Rebound Relays and Inhibitory Vetoes Stabilize Sparse Sequential Activity in HVC": Methods

We constructed single-compartment Hodgkin-Huxley type models of HVC neurons, incorporating pharmacologically validated ionic and synaptic currents[1–5]. Model neuron of $HVC_{RA}$, $HVC_X$, and $HVC_{INT}$ neurons were coupled via biologically realistic synapses and simulated using MATLAB's ode45 solver. Activation/inactivation kinetics and time constants were adapted from established neuronal models[1,6–9], with subtype-specific variation restricted to maximal ionic and synaptic conductances. Source code for all networks will be made publicly available through our lab website and ModelDB.

Ion channels of model HVC neurons

The mathematical equations developed for this network architecture that govern the voltage dynamics for each class of HVC neurons are described in detail in Bou Diab et al[10]. In short, each model neuron in the network is a single-compartment model of the Hodgkin-Huxley type with ionic currents integrated based on the class of the neuron and the underlying ionic currents that are expressed in it pharmacologically[1–4]. For example, $HVC_X$ and $HVC_{INT}$ model neurons exhibit the T-type $Ca^{++}$ and hyperpolarization-activated inward currents, $HVC_X$ and $HVC_{RA}$ neurons express the $Ca^{++}$ - dependent $K^+$ current, $HVC_{RA}$ neurons exhibit the A-type $K^+$ current, while all three classes express the fast-spiking $Na^+$ and the delayed rectifier $K^+$ currents[1]. The membrane potential of each HVC neuron obeys the following equations:

$$C_m \frac{dV_{RA}}{dt} = -I_L - I_K - I_{Na} - I_{CaL} - I_A - I_{SK} \quad (1)$$

$$C_m \frac{dV_X}{dt} = -I_L - I_K - I_{Na} - I_{CaL} - I_{CaT} - I_{SK} - I_H \quad (2)$$

$$C_m \frac{dV_{INT}}{dt} = -I_L - I_K - I_{Na} - I_{CaL} - I_{CaT} - I_H \quad (3)$$

where $C_m$ is the membrane capacitance. The associated equations and parameters for each of the activation/inactivation gating variables for each ionic current are given in Daou et al.[1] and shown below (Table 1). In total, every single model $HVC_{RA}$, $HVC_X$, and $HVC_{INT}$ neuron had a total of 6, 8 and 7 ODEs, respectively, that govern their intrinsic dynamics. Every synaptic current that was integrated to any model neuron added a new ODE to the set of ODEs governing the membrane potential of the corresponding model neuron.

### *Voltage-gated ionic currents*

The constant-conductance leak current is $I_L = g_L(V - V_L)$. The remaining voltage-gated ionic currents have non-constant dependent currents with activation/inactivation kinetics:

$$I_K = g_K n^4 (V - V_K) \quad (4)$$

$$I_{Na} = g_{Na} m_\infty^3(V) h (V - V_{Na}) \quad (5)$$

$$I_A = g_A a_\infty(V) e (V - V_K) \quad (6)$$

$$I_{CaL} = g_{Ca} s_\infty^2(V) \frac{Ca_{ex}}{1 - e^{\frac{2FV}{RT}}} \quad (7)$$

where $x_\infty(V) = \frac{1}{1 + e^{\frac{V - \theta_x}{\sigma_x}}}$ , $\quad x = m, a \text{ or } s \quad$ (8)

and $\frac{dx}{dt} = \frac{x_\infty(V) - V}{\tau_x}$ , $x = n, h \text{ or } e \quad$ (9)

where $x_\infty(V)$ for n and e is given by (8) and for $h_\infty$ as follows

$$h_\infty(V) = \frac{\alpha_h(V)}{\alpha_h(V) + \beta_h(V)} \quad (10)$$

where $\alpha_h(V) = 0.128e^{\frac{V+15}{-18}}$ (11)

and $\beta_h(V) = \frac{4}{1+e^{\frac{V+27}{-5}}}$ (12)

### *Low-voltage activated T-type calcium current*

The low-voltage activated T-type $Ca^{2+}$ current is described by the Goldman-Hodgkin-Katz formula:

$I_{CaT} = g_{CaT}\,(a_T)^3_\infty(V)(b_T)^2_\infty(r_T)\frac{Ca_{ex}}{1-e^{\frac{2FV}{RT}}}$ (13)

where $a_{T\infty}(V) = \frac{1}{1+e^{\frac{V-\theta_{a_T}}{\sigma_{a_T}}}}$ (14)

and $b_{T\infty}(r_T) = \frac{1}{1+e^{\frac{r_T-\theta_b}{\sigma_b}}} - \frac{1}{1+e^{\frac{-\theta_b}{\sigma_b}}}$ (15)

with $\frac{dr_T}{dt} = \frac{r_{T\infty}(V)-r_T}{\tau_{r_T}(V)}$ (16)

and $r_{T\infty}(V) = \frac{1}{1+e^{\frac{V-\theta_{a_T}}{\sigma_{a_T}}}}$ (17)

and $\tau_{r_T}(V) = \tau_{r_0} + \frac{\tau_{r_1}}{1+e^{\frac{V-\theta_{r_{r_T}}}{\sigma_{r_{r_T}}}}}$ (18)

### *Calcium-dependent potassium current*

The small conductance potassium current ($I_{SK}$ ) is modeled as

$I_{SK} = g_{SK}\,k_\infty([Ca^{2+}]_i)(V-V_K)$ (19)

where $k_\infty([Ca^{2+}]_i) = \frac{[Ca^{2+}]_i^2}{[Ca^{2+}]_i^2+k_s^2}$ (20)

and $\frac{d[Ca^{2+}]_i}{dt} = -f\,\varepsilon(I_{CaL} + I_{CaT}) + k_{Ca}([Ca^{2+}]_i - b_{Ca})$ (21)

#### *Hyperpolarization-activated inward current*

The hyperpolarization activated inward current's activation is modeled as in Destexhe and Babloyantz[6] using a fast component $(r_f)$ and a slow component $(r_s)$ as follows:

$$I_H = g_H\,[k_r r_f + (1 - k_r)\, r_s](V - V_h) \quad (22)$$

The fast activation component is given by:

$$\frac{dr_f}{dt} = \frac{r_{f\infty}(V) - r_f}{\tau_{r_f}(V)} \quad (23)$$

where $r_{f_\infty}(V) = \frac{1}{1+e^{\frac{V-\theta_{r_f}}{\sigma_{r_f}}}}$ (24)

with its time constant $\tau_{r_f}(V) = \frac{p_{r_f}}{\frac{-7.4(V+70)}{e^{(\frac{V+70}{-0.8})}-1}+65\,e^{(\frac{V+56}{-23})}}$ (25)

The slow activation component obeys:

$$\frac{dr_s}{ds} = \frac{r_{s\infty}(V) - r_s}{\tau_{r_s}} \quad (26)$$

where $r_{s_\infty}(V) = \frac{1}{1+e^{\frac{-(V-\theta_{r_s})}{\sigma_{r_s}}}}$ (27)

| Parameter | Value | Parameter | Value |
|---|---|---|---|
| $\boldsymbol{V_L}$ | $-70\ mV$ | $\theta_{a_T}$ | $-65\ mV$ |
| $\boldsymbol{V_K}$ | $-90\ mV$ | $\theta_{\boldsymbol{b}}$ | $0.4\ mV$ |
| $\boldsymbol{V_{Na}}$ | $50\ mV$ | $\theta_{r_T}$ | $-67\ mV$ |
| $\boldsymbol{V_{Ca}}$ | $85\ mV$ | $\theta_{r_{r_T}}$ | $68\ mV$ |
| $\boldsymbol{V_H}$ | $-30\ mV$ | $\boldsymbol{\sigma_m}$ | $-5\ mV$ |
| $\boldsymbol{g_L}$ | $2\ nS$ | $\boldsymbol{\sigma_n}$ | $-5\ mV$ |
| $\boldsymbol{g_{Ca}}$ | $19\ nS$ | $\boldsymbol{\sigma_s}$ | $-0.05\ mV$ |
| $\boldsymbol{\tau_n}$ | $10\ msec$ | $\boldsymbol{\sigma_a}$ | $-10\ mV$ |
| $\boldsymbol{\tau_{hp}}$ | $1000\ msec$ | $\boldsymbol{\sigma_e}$ | $5\ mV$ |
| $\boldsymbol{\tau_e}$ | $20\ msec$ | $\boldsymbol{\sigma_{r_f}}$ | $5\ mV$ |
| $\boldsymbol{\tau_h}$ | $1\ msec$ | $\boldsymbol{\sigma_{r_s}}$ | $25\ mV$ |
| $\boldsymbol{\tau_{r_s}}$ | $1500\ msec$ | $\boldsymbol{\sigma_{a_T}}$ | $-7.8\ mV$ |
| $\boldsymbol{\tau_{r_0}}$ | $200\ msec$ | $\boldsymbol{\sigma_b}$ | $-0.1\ mV$ |
| $\boldsymbol{\tau_{r_1}}$ | $87.5\ msec$ | $\boldsymbol{\sigma_{r_T}}$ | $2\ mV$ |
| $\boldsymbol{\theta_m}$ | $-35\ mV$ | $\boldsymbol{\sigma_{r_{r_T}}}$ | $2.2\ mV$ |
| $\boldsymbol{\theta_n}$ | $-30\ mV$ | $f$ | 0.1 |
| $\boldsymbol{\theta_s}$ | $-20\ mV$ | $\boldsymbol{\varepsilon}$ | $0.0015\ pA^{-1}$ microM $msec^{-1}$ |
| $\boldsymbol{\theta_a}$ | $-20\ mV$ | $\boldsymbol{k_{Ca}}$ | $0.3\ msec^{-1}$ |
| $\boldsymbol{\theta_e}$ | $-60\ mV$ | $\boldsymbol{b_{Ca}}$ | $0.1\ micro\ M$ |
| $\boldsymbol{\theta_{r_f}}$ | $-105\ mV$ | $\boldsymbol{k_s}$ | $0.5\ micro\ M$ |

| | | | |
|---|---|---|---|
| $\boldsymbol{\theta_{r_s}}$ | $-105\ mV$ | $\boldsymbol{p_{r_f}}$ | 100 |

Synaptic currents

The three classes of HVC neurons were connected via synaptic currents that were discovered pharmacologically[4]. Model $HVC_{RA}$ and $HVC_X$ neurons excite $HVC_{INT}$ neurons via AMPA currents, while $HVC_{INT}$ neurons inhibits both classes of projections neurons via $GABA_A$ current[10]. Each synaptic current represents the synaptic input(s) from the presynaptic cell(s) to the particular HVC model neuron, and is modeled as $I_{syn} = \sum_X I_{X\to Y}$ where $I_{X\to Y} = g_{X\to Y} s_{X\to Y}(V - V_{X\to Y})$. Here the summation is taken over the presynaptic HVC neurons where X represents a presynaptic cell, Y represents a postsynaptic cell, $V_{X\to Y}$ is the reversal potential for the synapse in the postsynaptic cell with $V_{X\to Y} = V_{AMPA}$ for excitatory input and $V_{X\to Y} = V_{GABA-A}$ for inhibitory input.

The model equations for the synaptic currents are shown below, taken after [11,12].

$I_{AMPA} = \overline{g_{AMPA}} s_{AMPA}(V - V_{AMPA})$ (28)
$I_{GABA-A} = \overline{g_{GABA-A}} s_{GABAA}(V - V_{GABA-A})$ (29)

where $s_{AMPA}$ and $s_{GABAA}$ are given by

$[T]_{pre} = \frac{T_{max}}{1+\exp(\frac{V_{pre}-V_T}{K_p})}$ and $\frac{d_s}{d_t} = a_r\,[T](1-s) - \ a_d s$ (30)

with $T_{max} = 1, K_p = 5, V_T = 2$. For $GABA_A$, $a_r = 5$ and $a_d = 0.18$, while for AMPA, $a_r = 1.1$ and $a_d = 0.19$.

Synaptic interactions in all networks were restricted to AMPA-mediated excitation and $GABA_A$-mediated inhibition. We excluded NMDA and $GABA_B$ currents because their complex dependencies (e.g., $Mg^{2+}$ block, G-protein signaling) require additional parameters and differential equations that remain uncharacterized in HVC, and because they are not essential for capturing the excitation-inhibition balance underlying our network dynamics. In the model, $HVC_{RA}$ neurons provide AMPA excitation to $HVC_{RA}$ and $HVC_{INT}$ neurons; $HVC_{INT}$ neurons deliver $GABA_A$ inhibition to $HVC_{RA}$ and $HVC_X$ neurons; and $HVC_X$ neurons excite $HVC_{INT}$ and $HVC_{RA}$ neurons via AMPA inputs.

Moreover, we reproduced the voltage traces elicited by the dual intracellular recordings conducted by Mooney and Prather[4] using dual intracellular recordings by tuning the synaptic parameters (excitatory and inhibitory currents' activation/inactivation constants, etc..) to ensure that our excitatory and synaptic currents in the model can replicate intricate features of synaptic transmission, like strengths of excitation/inhibition, magnitudes of voltage deflections and other trace morphologies[10]. We did so by connecting each two HVC neurons of different classes together using the corresponding synaptic currents (AMPA or GABA), and fitting the model voltage traces elicited to their corresponding traces shown by Mooney and Prather[4] using their dual-intracellular recording studies (Figure 3 in Bou Diab et al[10]). In short, DC-evoked action potentials in any of the HVC neuronal classes trigger excitatory or inhibitory postsynaptic potentials in their postsynaptic counterpart based on the synaptic currents expressed[2,4]. This exercise was considered as a basis to our network modeling that will be described next. The equations and associated ODEs for the synaptic currents, as well as the intrinsic and synaptic parameters that govern the kinetics of the activation/inactivation variables of all ionic and synaptic currents are described in Bou Diab et al[10].

Parameter variations and model assessment criteria

The main aim behind the network architecture is to generate realistic *in vivo* firing activity as seen by the three classes of HVC neurons during singing. The focus is on 1) the accurate propagation of sequential activity orchestrated by the various microcircuits in the network and 2) the generation of biologically realistic spiking/bursting patterns for each neuronal class as observed through the *in vivo* intracellular recordings conducted[3,13,14]. This includes maintaining spike shapes, spike frequency, sags and/or rebound upon inhibition, resting membrane potential, burst patterns, rebound firing/bursting, subthreshold oscillations, and other intrinsic properties that the different classes of HVC neurons exhibit[1,5]. In short, the network behavior was considered optimal if the model voltage traces for each class matched the following criteria:

1. $HVC_{RA}$ neurons exhibit time-locked characteristic bursting (a single burst with 3-6 spikes within ~10 ms), with spikes riding on a plateau like what's seen during singing.
2. $HVC_{X}$ neurons produce 1-4 bursts (4-9 spikes each burst), which are time-locked and primarily rebound bursts following inhibition[15,16].
3. $HVC_{INT}$ neurons show tonic activation with continuous spiking and bursting throughout the song.

Automated adjustment of model parameters was performed to qualitatively reproduce desired membrane potential trajectories, as described next. All parameters that govern the voltage dynamics for the three neuronal subtypes were fixed in our model (Table 1), similar to what we did in Bou Diab et al[10], and the only parameters that were allowed to vary were a subset of the maximal conductances of synaptic and intrinsic currents that are shown in Supplementary Figure 1. Within the population of ionic maximal conductances, the $Na^+$ conductance ($g_{Na}$), the $K^+$ conductance ($g_K$) and high-threshold L-type $Ca^{++}$ conductance ($g_{CaL}$) were fixed to values that realistically fit the spike morphologies (upstrokes and downstrokes of action potentials, plateaus, etc…) for each class of HVC neurons, in response to a depolarizing applied current pulse given *in vitro*[1,10]. Moreover, the effects of varying the A-type $K^+$ conductance ($g_A$) and the $Ca^{++}$ -dependent $K^+$ conductance ($g_{SK}$) on $HVC_{RA}$ neurons' intrinsic properties as well as on the propagation of sequential activity had been described earlier in detail[10], so we fixed them in this network to values that generate realistic bursting patterns, rather than repeating the same exercise again.

The intrinsic ionic maximal conductances that were randomly varied and that played key roles in orchestrating the network dynamics were the following: 1) $Ca^{++}$ dependent $K^+$ conductance ($g_{SK}$) for $HVC_X$ neurons and 2) hyperpolarization-activated inward conductance ($g_H$) for $HVC_X$ and $HVC_{INT}$ neurons and 3) low-threshold T-type $Ca^{++}$ conductance ($g_{CaT}$) for $HVC_X$ and $HVC_{INT}$ neurons (Supplementary Figure 1A). Unlike intrinsic maximal conductances, all synaptic maximal conductances were randomly varied (Supplementary Figure 1B).

Random variations in every parameter (intrinsic and synaptic) that was allowed to vary were conducted to qualitatively reproduce the membrane potential trajectories of the three classes of HVC neurons as observed during the bird's singing and described in Bou Diab et al[10]. In short, every varying parameter was randomly increased up or decreased down by simulating the network many times, until the point that the network activity ceased to generate the optimal behavior as described earlier, and thereby delineating the legitimate ranges of variation. This approach also gave us a measure of the network robustness to varied parameter values.

Moreover, and similar to what was described earlier[10], we conducted the variations at the population level as well as the individual neuronal level by either 1) setting the maximal conductance for all neurons of the same class to a single value and randomly changing that value for all, or 2) varying the maximal conductance for one neuron at a time of the same class while fixing the others to their default values. If any of the simulations generated voltage traces (for any of the neurons in the network) that did not adhere to the desired network behavior criteria described earlier at the levels of intrinsic and synaptic properties, then the corresponding parameters combination is ignored (more details on that are described in Bou Diab et al[10]). The constraints that we added, though stringent, ensured that we are reproducing biophysically realistic firing patterns for all classes of HVC neurons, and for each neuron within a particular class. The ranges that each of the synaptic and ionic maximal conductances that were allowed to vary are shown in Supplementary Figure 1.

Similar to what was done with the maximal ionic conductances, all synaptic conductances were systematically varied rather than fixed. Supplementary Figure 1B summarizes the ranges that preserved robust sequence propagation and in vivo-like activity across all neuronal classes. Most synaptic parameters tolerated broad variability, with the notable exception of the excitatory conductance from $HVC_X$ to $HVC_{RA}$. Increasing this synaptic conductance to larger magnitudes would induce a strong excitation in $HVC_{RA}$ generating non-realistic bursting in their spike train[3].

<u>Extrinsic input to HVC neurons</u>

None of the neurons in our network receive external input from outside the HVC. The network is kick-started by a stochastic DC input to the first $HVC_{RA}$ neuron of the very first microcircuit, that is to $HVC_{RA_1}^{1}$. In other words, $HVC_{RA_1}^{1}$ is the only projection neuron in this network that receives input from outside HVC. The stochastic input current is of the form $I_{noise}(t) = \sigma.\xi(t)$ where $\xi(t)$ is a Gaussian white noise with zero mean and unit variance, and $\sigma$ is the noise amplitude. To make HVC tonic bursting interneurons spike densely throughout the song, we injected every tonic bursting interneuron with a stochastic input. The stochastic current is added in the differential equation dictating the membrane potential of the model interneuron. This stochastic drive mimics the fluctuations in synaptic input arising from random presynaptic activity and background noise. For values of $\sigma$ within 1-5% of the mean synaptic conductance, the stochastic current has no effect on network propagation. For larger values of $\sigma$, the desired network activity was disrupted or halted. The ability of the model networks to reproduce the sequential propagation in the presence of this stochastic input is a strong indicator of the robustness of our model. The propagation of sequential activity along with the realistic firing of the three classes of HVC neurons is maintained and orchestrated by HVC's intrinsic and synaptic processes without relying on extrinsic inputs.

### References


1. Daou, A., Ross, M.T., Johnson, F., Hyson, R.L., and Bertram, R. (2013). Electrophysiological characterization and computational models of HVC neurons in the zebra finch. J Neurophysiol *110*, 1227–1245. https://doi.org/10.1152/jn.00162.2013.

2. Kosche, G., Vallentin, D., and Long, M.A. (2015). Interplay of inhibition and excitation shapes a premotor neural sequence. J Neurosci *35*, 1217–1227. https://doi.org/10.1523/JNEUROSCI.4346-14.2015.

3. Long, M.A., Jin, D.Z., and Fee, M.S. (2010). Support for a synaptic chain model of neuronal sequence generation. Nature *468*, 394–399. https://doi.org/10.1038/nature09514.

4. Mooney, R., and Prather, J.F. (2005). The HVC microcircuit: the synaptic basis for interactions between song motor and vocal plasticity pathways. J Neurosci *25*, 1952–1964. https://doi.org/10.1523/JNEUROSCI.3726-04.2005.

5. Daou, A., and Margoliash, D. (2020). Intrinsic neuronal properties represent song and error in zebra finch vocal learning. Nat Commun *11*, 952. https://doi.org/10.1038/s41467-020-14738-7.

6. Destexhe, A., and Babloyantz, A. (1993). A model of the inward current Ih and its possible role in thalamocortical oscillations. Neuroreport *4*, 223–226. https://doi.org/10.1097/00001756-199302000-00028.

7. Dunmyre, J.R. (2011). Qualitative validation of the reduction from two reciprocally coupled neurons to one self-coupled neuron in a respiratory network model. J Biol Phys *37*, 307–316. https://doi.org/10.1007/s10867-011-9213-0.

8. HODGKIN, A.L., and HUXLEY, A.F. (1952). A quantitative description of membrane current and its application to conduction and excitation in nerve. J Physiol *117*, 500–544. https://doi.org/10.1113/jphysiol.1952.sp004764.

9. Terman, D., Rubin, J.E., Yew, A.C., and Wilson, C.J. (2002). Activity patterns in a model for the subthalamopallidal network of the basal ganglia. J Neurosci *22*, 2963–2976. https://doi.org/10.1523/JNEUROSCI.22-07-02963.2002.

10. Bou Diab, Z., Chammas, M., and Daou, A. (2025). Biophysical network modeling of temporal and stereotyped sequence propagation of neural activity in the premotor nucleus HVC. Elife *14*. https://doi.org/10.7554/eLife.105526.

11. Destexhe, A., Mainen, Z.F., and Sejnowski, T.J. (1994). Synthesis of models for excitable membranes, synaptic transmission and neuromodulation using a common kinetic formalism. J Comput Neurosci *1*, 195–230. https://doi.org/10.1007/BF00961734.

12. Varela, J.A., Sen, K., Gibson, J., Fost, J., Abbott, L.F., and Nelson, S.B. (1997). A quantitative description of short-term plasticity at excitatory synapses in layer 2/3 of rat primary visual cortex. J Neurosci *17*, 7926–7940. https://doi.org/10.1523/JNEUROSCI.17-20-07926.1997.

13. Hahnloser, R.H.R., Kozhevnikov, A.A., and Fee, M.S. (2002). An ultra-sparse code underlies the generation of neural sequences in a songbird. Nature *419*, 65–70. https://doi.org/10.1038/nature00974.

14. Kozhevnikov, A.A., and Fee, M.S. (2007). Singing-related activity of identified HVC neurons in the zebra finch. J Neurophysiol *97*, 4271–4283. https://doi.org/10.1152/jn.00952.2006.

15. Lewicki, M.S. (1996). Intracellular characterization of song-specific neurons in the zebra finch auditory forebrain. J Neurosci *16*, 5855–5863.

16. Mooney, R. (2000). Different subthreshold mechanisms underlie song selectivity in identified HVc neurons of the zebra finch. J Neurosci *20*, 5420–5436. https://doi.org/10.1523/JNEUROSCI.20-14-05420.2000.
