## Supplementary Figures for "Rebound Relays and Inhibitory Vetoes Stabilize Sparse Sequential Activity in HVC"

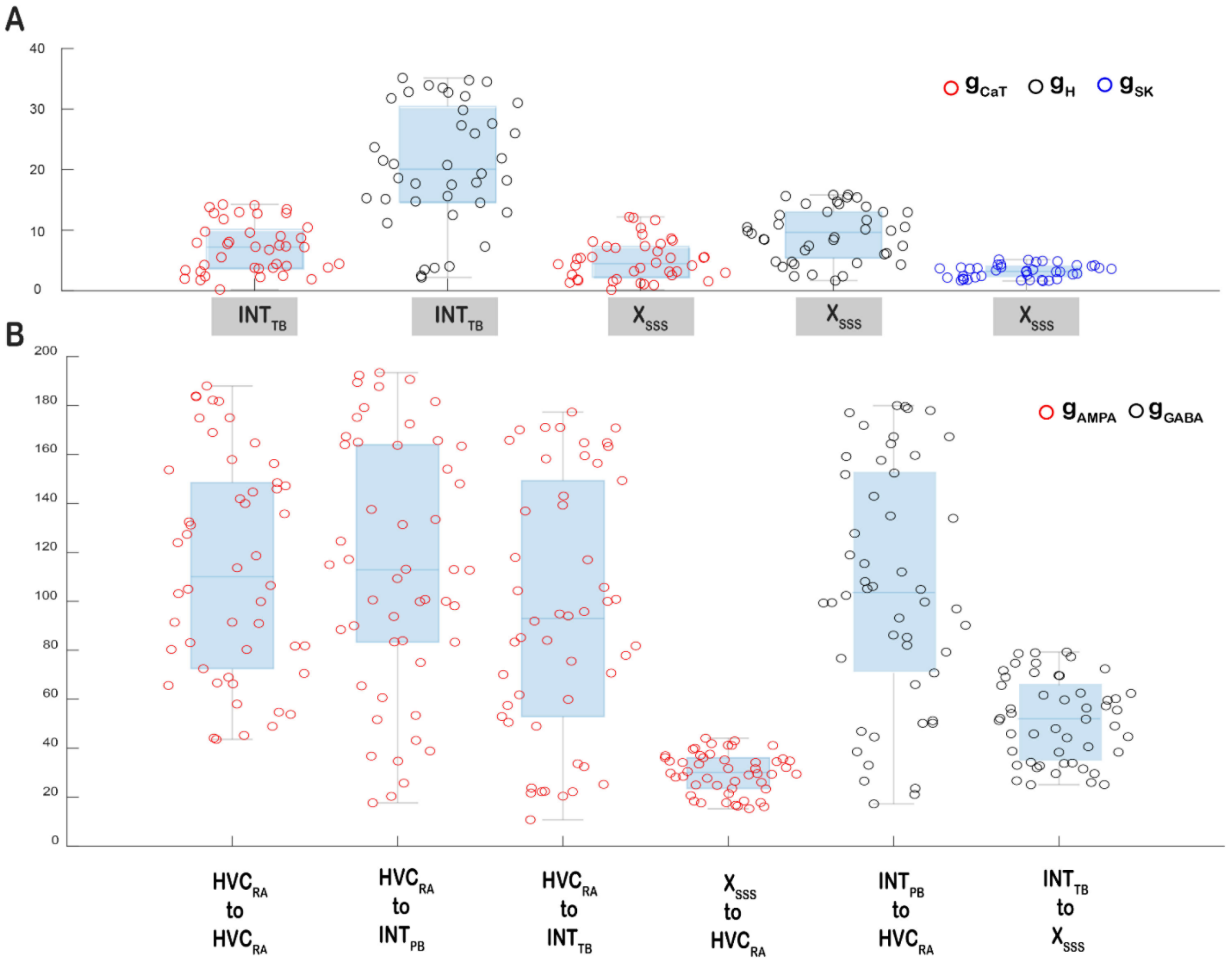


**Supplementary Figure 1: Parameter ranges for intrinsic and synaptic conductances in the HVC microcircuit model.** (**A**) Distributions of intrinsic maximal conductances controlling excitability and rebound in the microcircuit-specific tonic interneuron ($INT_{TB}$) and the microcircuit-specific X-projecting relay neuron ($X_{SSS}$): T-type $Ca^{2+}$ ($g_{CaT}$, red), H-current ($g_H$, black) and $Ca^{2+}$-activated $K^+$ ($g_{SK}$, blue; shown for $X_{SSS}$). (**B**) Distributions of synaptic conductances implementing the model's core motifs: AMPAergic coupling within the $HVC_{RA}$ chain ($HVC_{RA} \rightarrow HVC_{RA}$), excitation from $HVC_{RA}$ to phasic and tonic interneurons ($HVC_{RA} \rightarrow INT_{PB}$; $HVC_{RA} \rightarrow INT_{TB}$), the cross-microcircuit relay from $X_{SSS}$ to the chain-initiating $HVC_{RA}$ neuron ($X_{SSS} \rightarrow HVC_{RA}$), and inhibitory control of propagation ($INT_{PB} \rightarrow HVC_{RA}$; $INT_{PB} \rightarrow X_{SSS}$; $g_{GABA}$, black). Open circles denote individual parameter draws; boxplots show median and interquartile range (whiskers indicate full range). Together, these distributions define the "allowed" parameter envelope used for the main simulations; departures from these ranges in subsequent perturbations destabilize firing regimes and/or terminate sequence propagation.

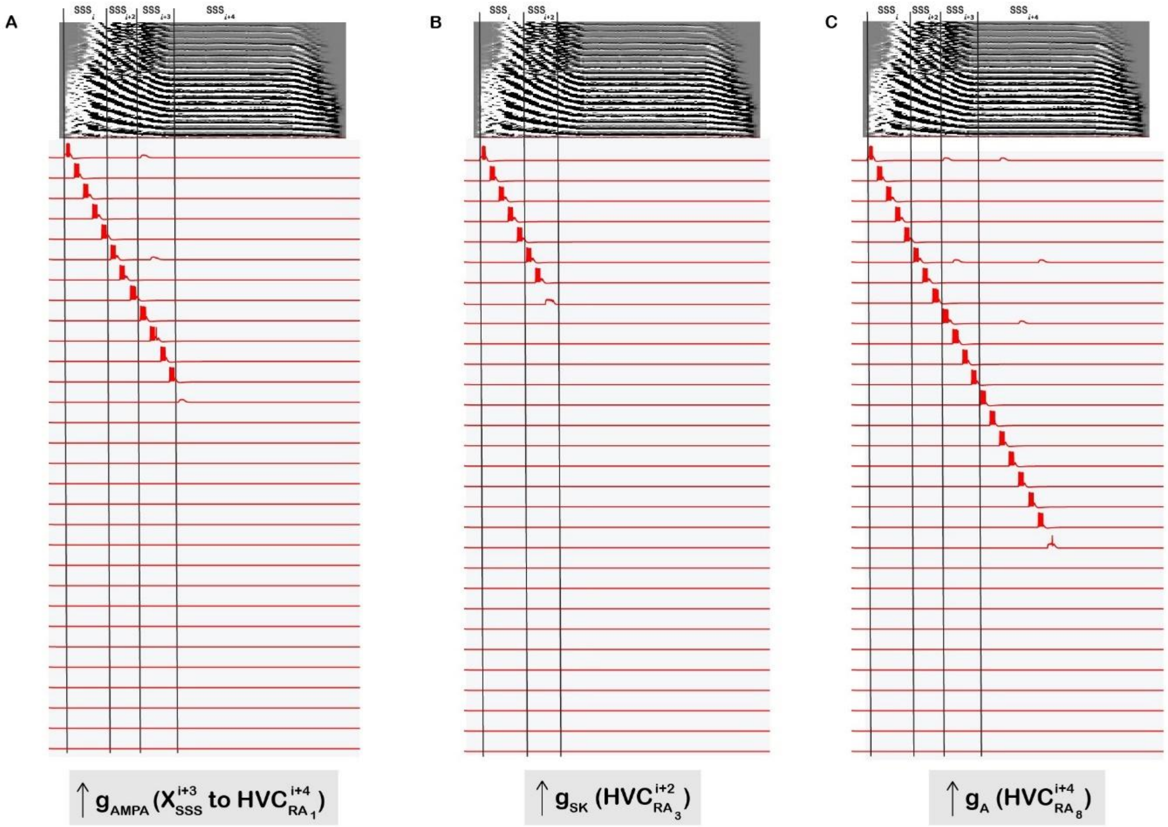


**Supplementary Figure 2: Intrinsic and synaptic parameter regimes that disrupt HVC$_{RA}$ bursting and halt sequence propagation.** Representative membrane potential traces from HVC$_{RA}$ neurons (red; subset shown) aligned to an example syllable segmented into sub-syllabic segments (SSSs; top waveform, vertical boundaries). Each panel illustrates a distinct mechanism by which the HVC$_{RA}$ chain fails, leading to premature termination of the sequence. (**A**) Increasing the AMPA coupling at the microcircuit relay ($g_{AMPA}$ from $X_{SSS}^{i+3}$ to the chain-initiating neuron $HVC_{RA_1}^{i+4}$) alters recruitment at the microcircuit boundary and can prevent reliable initiation of the next microcircuit when the effective relay drive falls outside the propagation-permissive range. (**B**) Increasing the $Ca^{2+}$-dependent $K^+$ conductance $g_{SK}$ in a representative HVC$_{RA}$ neuron (here $HVC_{RA_3}^{i+2}$) suppresses burst excitability, reducing spike output and ultimately eliminating the burst required for within-microcircuit propagation. (**C**) (C) Increasing the A-type $K^+$ conductance $g_A$ in a representative HVC$_{RA}$ neuron (here $HVC_{RA_8}^{i+4}$) delays and weakens recruitment, and when sufficiently large, abolishes HVCRA bursting, breaking the sequential chain. Together, these perturbations identify minimal synaptic gain (within-chain HVC$_{RA}$ → HVC$_{RA}$ and relay $X_{SSS} \rightarrow HVC_{RA_1}$) and bounded intrinsic "braking" ($g_A$, $g_{SK}$) as necessary conditions for maintaining ultra-sparse HVC$_{RA}$ bursting and uninterrupted motif-scale sequence propagation.

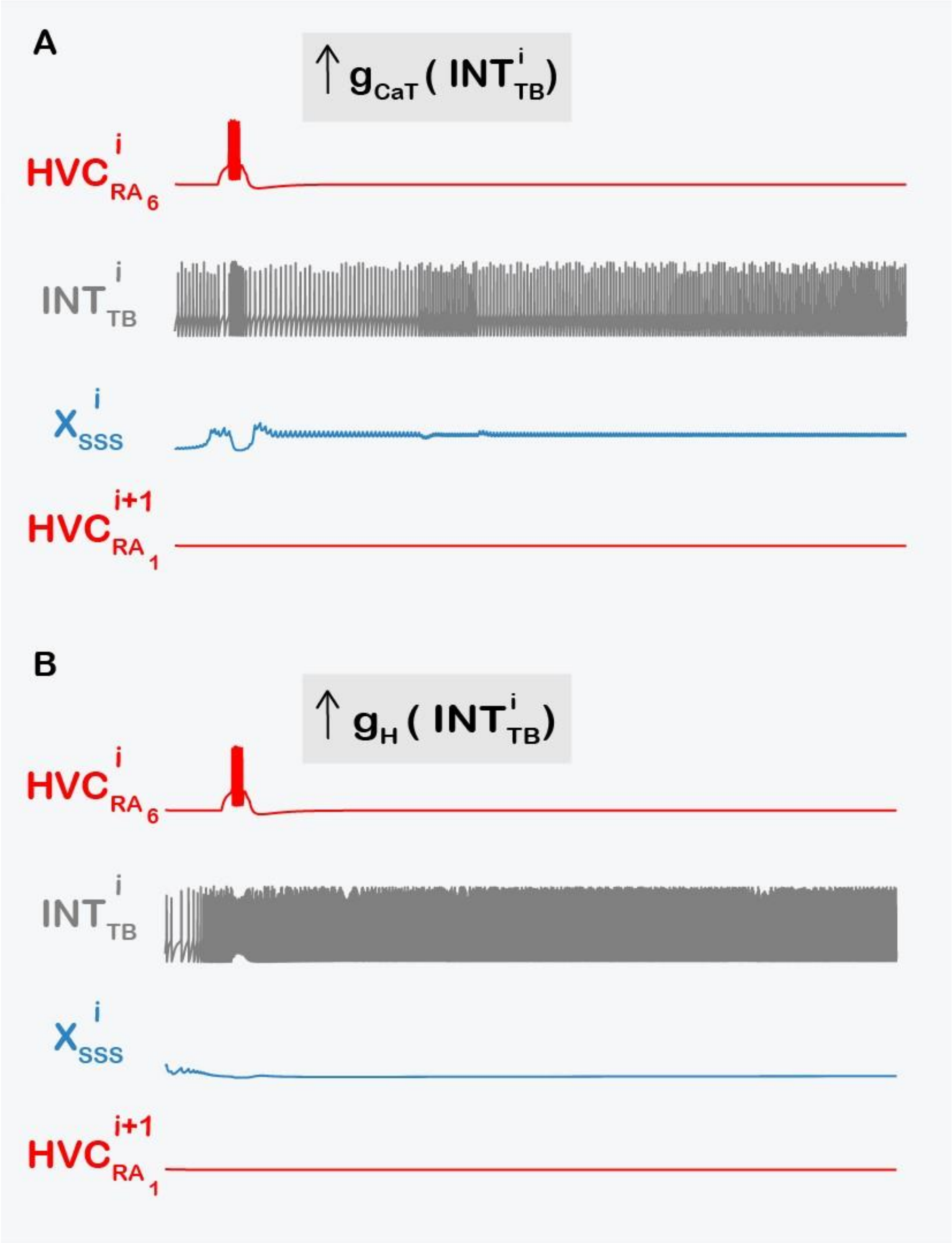


**Supplementary Figure 3: Runaway excitability of microcircuit-specific tonic interneurons clamps $X_{SSS}$ and halts sequence propagation.** Representative voltage traces illustrating failure modes in which increasing intrinsic rebound-related conductances in the microcircuit-specific tonic-bursting interneuron ($INT_{TB}^{i}$, gray) drives sustained interneuron activity and prevents rebound-mediated microcircuit transfer. In both panels, the terminal RA-projecting neuron in microcircuit $i$ ($HVC_{RA_6}^{i}$, red) initiates activity, but successful propagation requires that $INT_{TB}^{i}$ deliver temporally bounded inhibition to $X_{SSS}^{i}$ (blue) so that release can evoke a rebound burst capable of recruiting $HVC_{RA_1}^{i+1}$ (red). (**A**) Three-fold increase in $g_{CaT}$ in $INT_{TB}^{i}$ induces abnormally dense, near-continuous firing in the interneuron, producing prolonged inhibition of $X_{SSS}^{i}$ and suppressing rebound bursting; consequently, $HVC_{RA_1}^{i+1}$ is not recruited and the sequence terminates. (**B**) Five-fold increase in $g_H$ in $INT_{TB}^{i}$ similarly drives $INT_{TB}^{i}$ into a high-activity regime, effectively clamping $X_{SSS}^{i}$ and preventing the rebound burst needed to excite $HVC_{RA_1}^{i+1}$, thereby halting propagation at the microcircuit boundary.

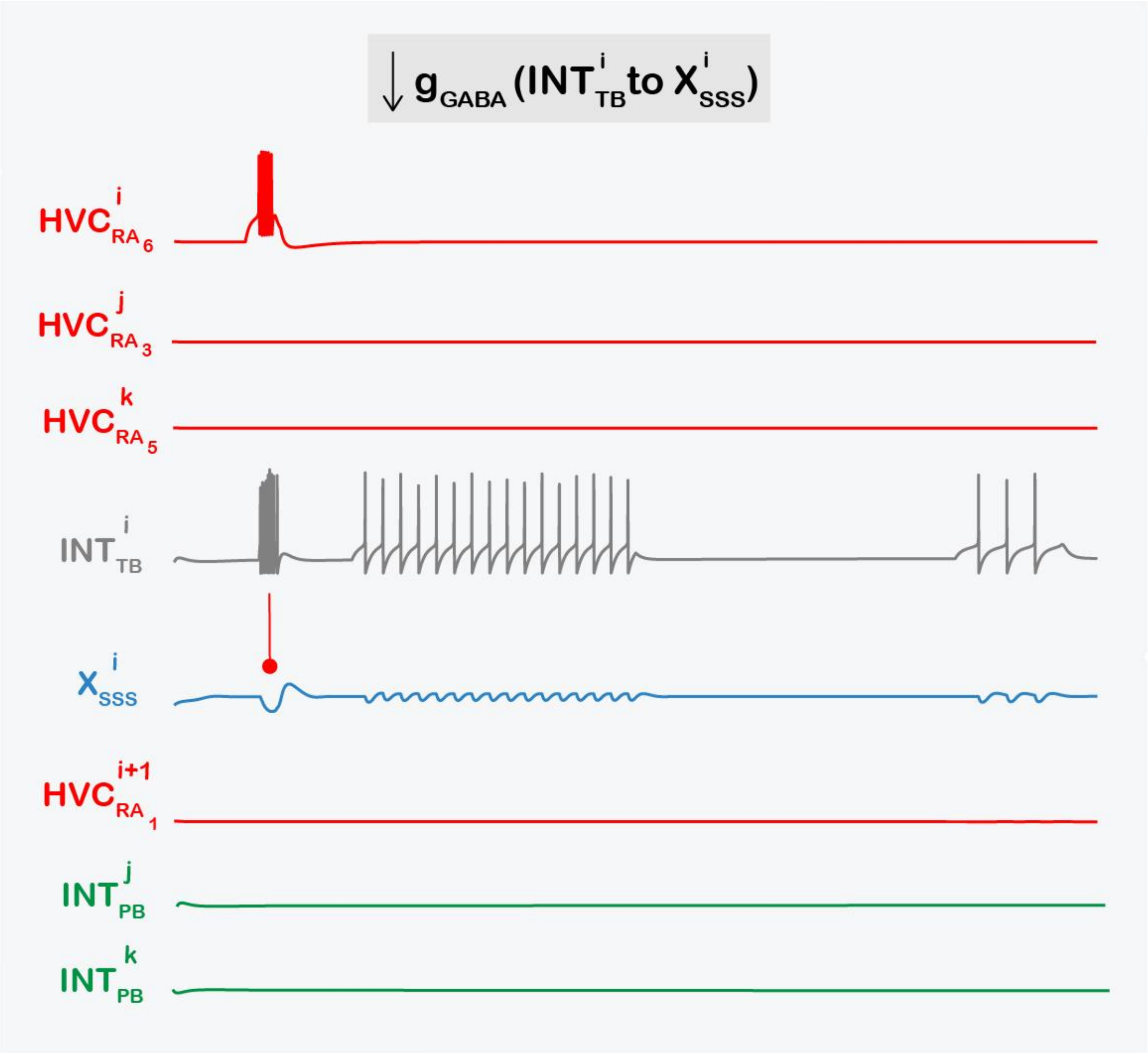


**Supplementary Figure 4: Weakening tonic inhibition onto microcircuit-specific HVCX abolishes rebound bursting and terminates sequence propagation.** Representative voltage traces showing a failure mode in which sequence propagation breaks when the inhibitory coupling from the microcircuit-specific tonic-bursting interneuron to its paired X-projecting neuron is reduced (10-fold decrease in $g_{GABA}$ from $INT_{TB}^{i}$ to $X_{SSS}^{i}$; label at top). The terminal RA-projecting neuron in microcircuit $i$ ($HVC_{RA_6}^{i}$, red) still excites $INT_{TB}^{i}$ (gray), eliciting a burst. This burst produces only a weak hyperpolarization in $X_{SSS}^{i}$ (blue; sag), which on release generates a small rebound overshoot rather than a suprathreshold rebound burst. Consequently, $X_{SSS}^{i}$ fails to recruit the chain-initiating neuron in the next microcircuit ($HVC_{RA_1}^{i+1}$, red), and propagation halts at the microcircuit boundary. Downstream HVCRA neurons ($HVC_{RA_3}^{j}$, $HVC_{RA_5}^{k}$) remain silent because activity never reaches their segments; accordingly, $INT_{TB}^{i}$ expresses only the initial burst and the phasic interneurons $INT_{PB}^{j}$ and $INT_{PB}^{k}$ (green) are not recruited due to the absence of HVCRA activity in microcircuits j and k.

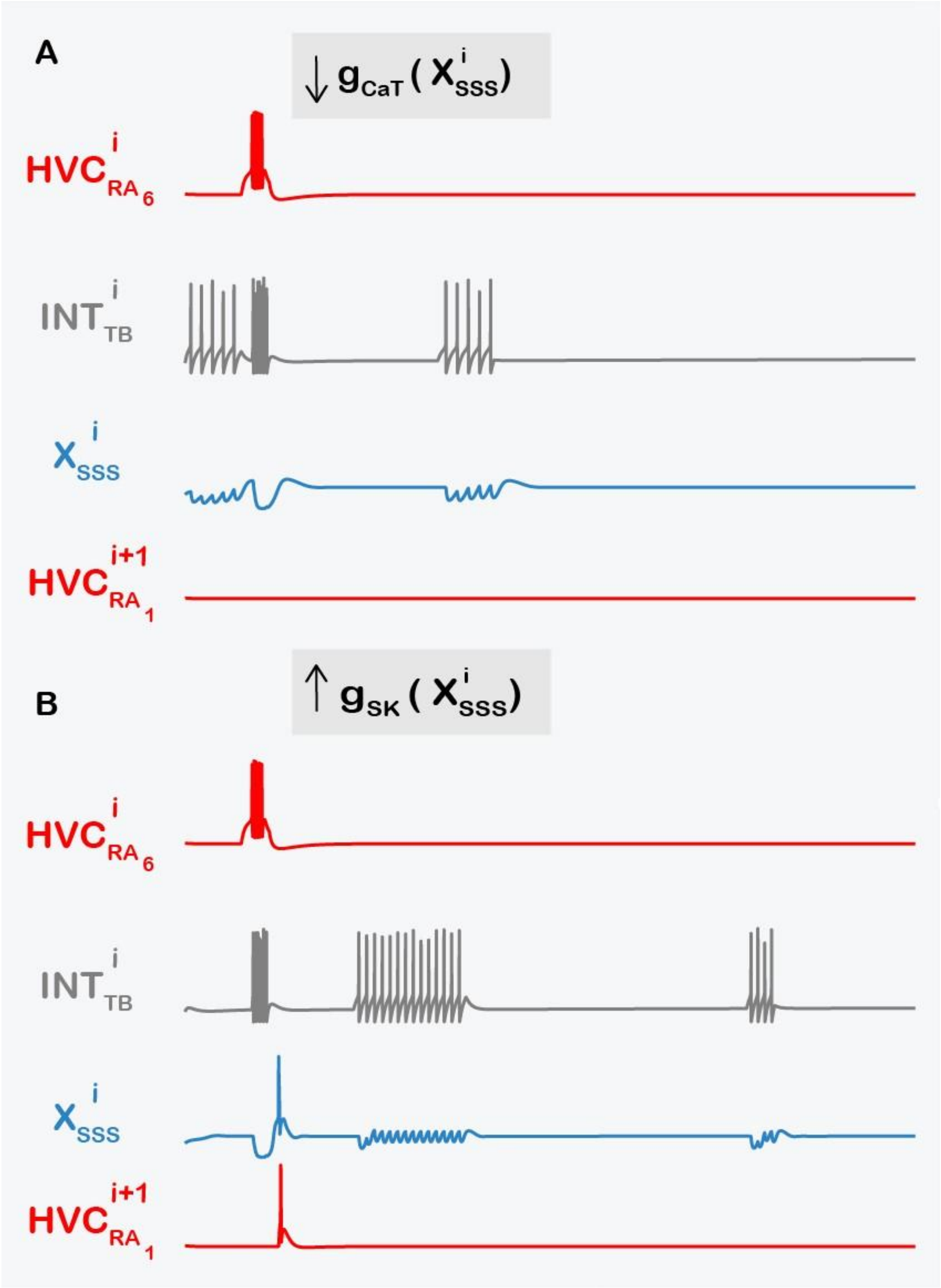


**Supplementary Figure 5: Intrinsic rebound excitability of $X_{SSS}$ gates microcircuit-to-microcircuit propagation.** Representative voltage traces illustrating how perturbing intrinsic conductances in the microcircuit-specific X-projecting neuron $X^{i}_{SSS}$ disrupts rebound bursting and halts sequence transfer to the next microcircuit. In both panels, the terminal RA-projecting neuron in microcircuit $i$ ($HVC^{i}_{RA_6}$, red) excites the microcircuit-specific tonic-bursting interneuron $INT^{i}_{TB}$ (gray), which inhibits $X^{i}_{SSS}$ (blue); successful propagation requires that release from this inhibition evokes a suprathreshold rebound burst in $X^{i}_{SSS}$ capable of recruiting the chain-initiating neuron $HVC^{i+1}_{RA_1}$ (red). (**A**) Reduced T-type $Ca^{2+}$ conductance (3-fold decrease in $g_{CaT}$ of $X^{i}_{SSS}$) weakens inward rebound drive, producing only subthreshold rebound depolarizations and preventing activation of $HVC^{i+1}_{RA_1}$, thereby terminating propagation at the microcircuit boundary. (**B**) Enhanced SK conductance (2-fold increase in $g_{SK}$ of $X^{i}_{SSS}$) potentiates the outward $Ca^{2+}$-dependent $K^{+}$ current ($I_{SK}$), counteracting rebound depolarization driven by $I_{CaT}$ (with support from $I_h$) and reducing or abolishing rebound spiking. As a consequence, the X → $HVC_{RA}$ relay fails and sequence propagation breaks.

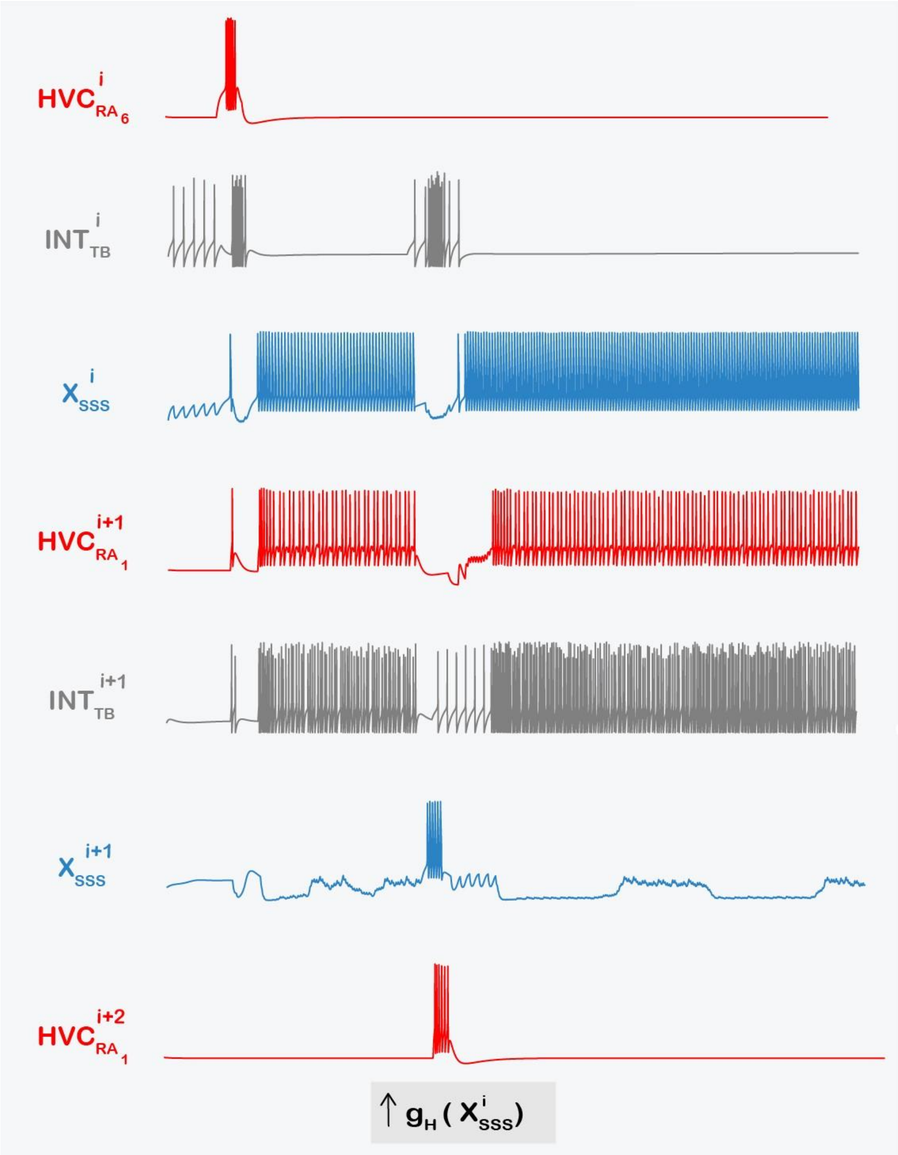


**Supplementary Figure 6: Excess $I_h$ in $X_{SSS}$ induces runaway excitation and disrupts song-like sequencing.** Voltage traces from successive microcircuits after a 5-fold increase in $g_H$ in the microcircuit-specific X-projecter $X_{SSS}^{i}$. Elevated $g_H$ drives $X_{SSS}^{i}$ into near-continuous firing, interrupted only during inhibition from $INT_{TB}^{i}$ (gray). This persistent activity propagates through the $X_{SSS}^{i} \rightarrow HVC_{RA_1}^{i+1}$ synapse, converting the normally sparse burst of $HVC_{RA_1}^{i+1}$ (red) into dense, continuous spiking and driving $INT_{TB}^{i+1}$ into high-frequency firing. The resulting strong inhibition largely clamps $X_{SSS}^{i+1}$, which produces only a single rebound burst during brief silent windows and recruits $HVC_{RA_1}^{i+2}$ at a markedly delayed time. Thus, sequence propagation becomes strongly phase-shifted and firing patterns become pathological ($HVC_{RA}$ dense spiking; $HVC_X$ sustained firing rather than 2-4 rebound bursts).

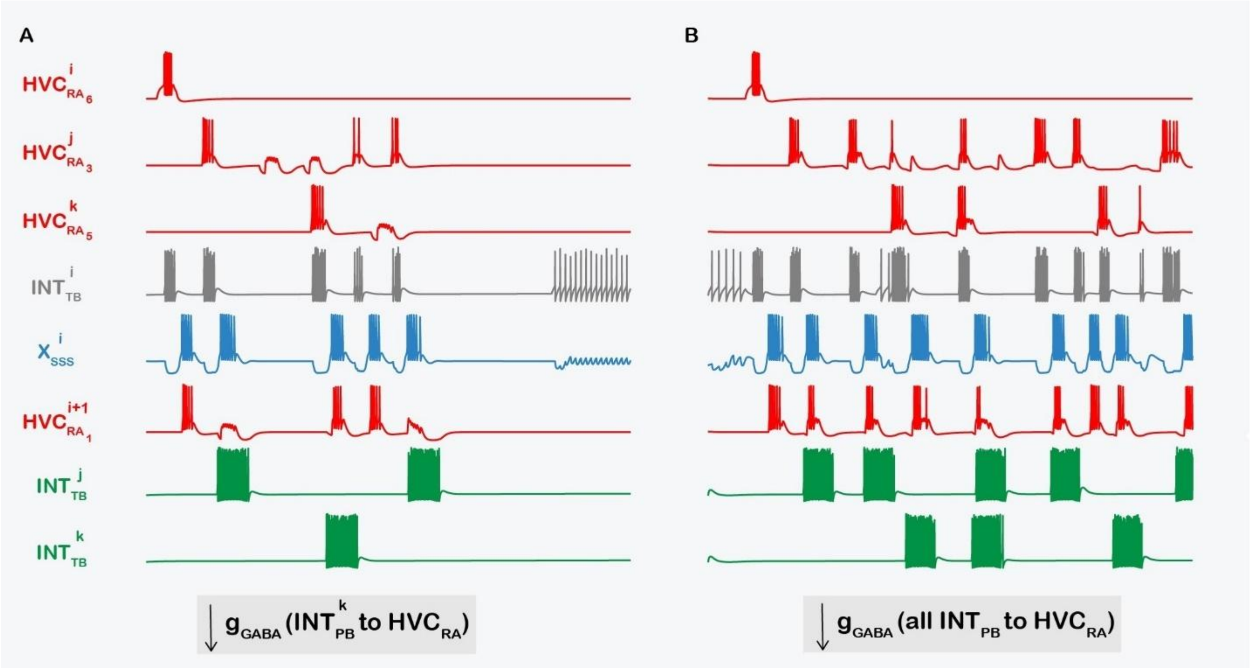


**Supplementary Figure 7: Weak phasic inhibition fails to veto off-time HVC$_{RA}$ bursts, triggering ectopic cascades and destabilizing the sequence.** Representative voltage traces illustrating how reducing GABAergic output from microcircuit-specific phasic-bursting interneurons ($INT_{PB}$, green) onto RA-projecting neurons ($HVC_{RA}$, red) corrupts song-like timing. In both panels, $X_{SSS}^{i}$ (blue) generates multiple rebound bursts, only the first of which should recruit the chain-initiating neuron $HVC_{RA_1}^{i+1}$; later rebounds would drive additional, wrong-time bursts unless suppressed by phasic "veto" inhibition. (**A**) *Localized perturbation*: $g_{GABA}$ from a single phasic burster ($INT_{PB}^{k}$) onto all of its HVC$_{RA}$ targets is reduced five-fold. $INT_{PB}^{j}$ (unperturbed) successfully suppresses the wrong-time burst during microcircuit $j$, whereas weakened $INT_{PB}^{k}$ fails to silence the later off-time burst. The resulting ectopic firing in $HVC_{RA_1}^{i+1}$ seeds additional downstream activity, producing a cascade of mistimed events through interneuron - X-projecter interactions. (B) *Global perturbation*: $g_{GABA}$ from all phasic bursters onto their HVC$_{RA}$ targets is reduced five-fold, broadly removing the temporal veto. This leads to pervasive off-schedule HVC$_{RA}$ bursting, increased interneuron recruitment, and widespread rebound-driven activation in X-projecters, yielding strongly non-physiological, destabilized sequence dynamics.

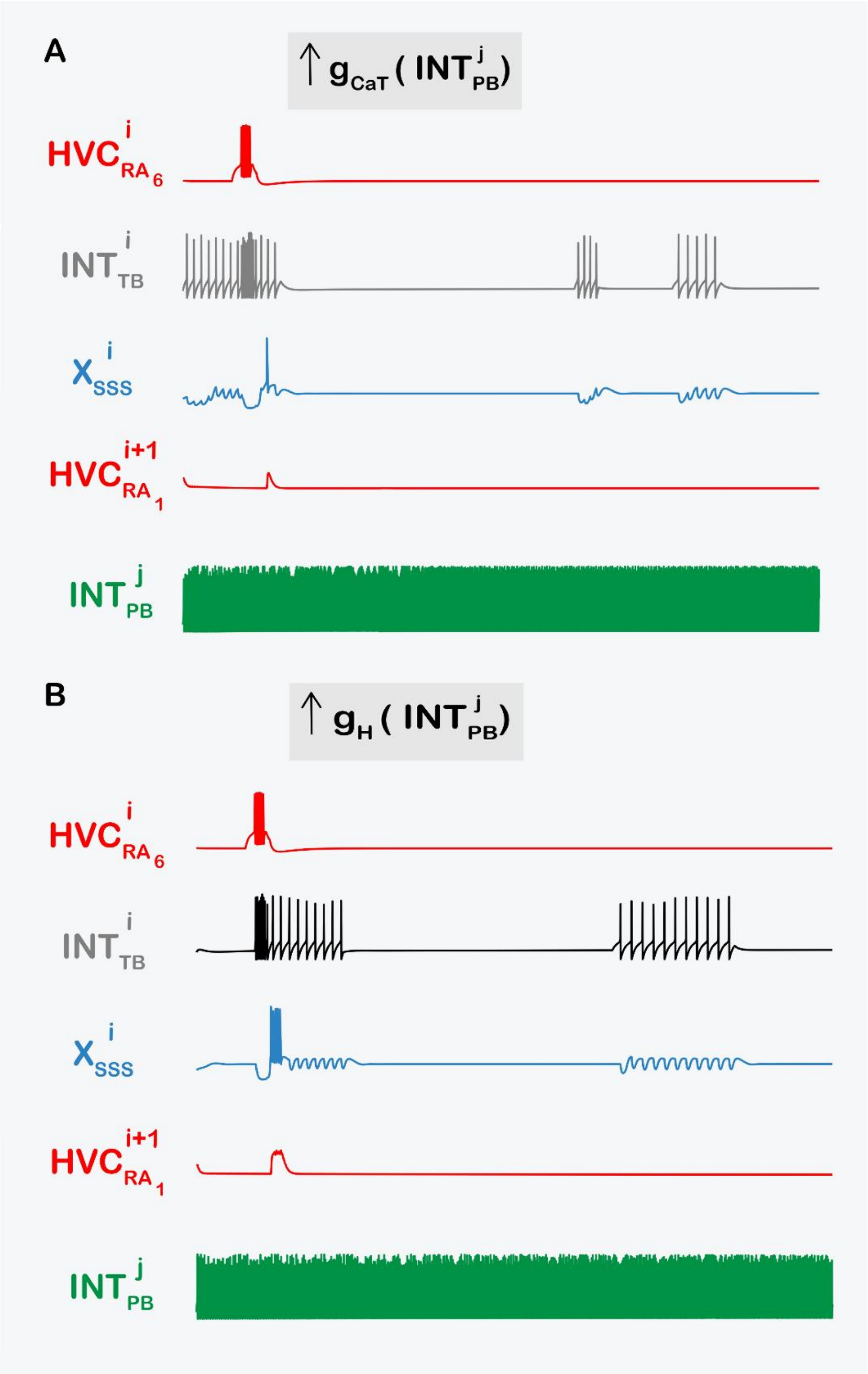


**Supplementary Figure 8: Upregulating $I_{CaT}$ or $I_h$ in a phasic burster converts segment-specific inhibition into a continuous clamp that silences HVCRA bursting**. Voltage traces showing that increasing intrinsic excitability of the microcircuit-specific phasic interneuron $INT_{PB}^{j}$ disrupts song-like dynamics by abolishing properly timed HVCRA bursts in its targets. Traces (top to bottom) include $HVC_{RA_6}^{i}$ (red), $INT_{TB}^{i}$ (gray), $X_{SSS}^{i}$ (blue), $HVC_{RA_1}^{i+1}$ (red), and the perturbed $INT_{PB}^{j}$ (green). (**A**) $g_{CaT}$ increased 4-fold. (**B**) $g_H$ increased 5-fold. In both cases, $INT_{PB}^{j}$ switches from phasic, SSS-locked activity to near-continuous firing, imposing persistent inhibition that prevents its connected HVCRA neurons from bursting.

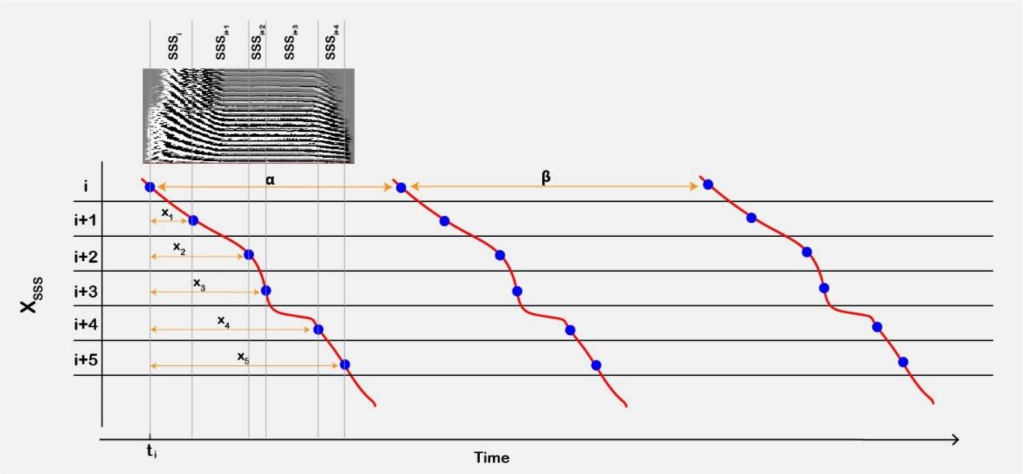


**Supplementary Figure 9: Schematic mechanism for parallel HVCX burst manifolds in the microcircuit chain.** Illustrative "neuron cascade" diagram for six successive microcircuit-specific X-projecting neurons ($X_{SSS}^{i}$ - $X_{SSS}^{i+5}$) showing how repeated rebound bursting can generate parallel burst manifolds across the population. For simplicity, each $X_{SSS}$ neuron is depicted as producing three bursts: an initial burst (blue) followed by two subsequent bursts (red), with fixed inter-burst intervals $\alpha$ (between the 1st and 2nd bursts) and $\beta$ (between the 2nd and 3rd bursts). Neurons are ordered by their initial burst times, which are offset by $x_1$, $x_2$, … relative to $t_i$. In the mode I, these offsets reflect propagation time through successive microcircuits and thus encode sub-syllabic segment (SSS) duration: longer SSSs recruit larger HVCRA ensembles and increase the delay between initial bursts of adjacent $X_{SSS}$ neurons. Subsequent bursts arise when tonic-bursting interneurons re-inhibit each $X_{SSS}$ neuron later in the motif, producing additional rebound events; preserving the same $\alpha$, $\beta$ structure across neurons yields quasi-linear manifolds of later bursts that are approximately parallel to the initial-burst manifold (compare to Figure 8B). Vertical lines and the overlaid example syllable indicate SSS boundaries within the canonical motif time base.
